# OneSC: A computational platform for recapitulating cell state transitions

**DOI:** 10.1101/2024.05.31.596831

**Authors:** Da Peng, Patrick Cahan

## Abstract

Computational modelling of cell state transitions has been a great interest of many in the field of developmental biology, cancer biology and cell fate engineering because it enables performing perturbation experiments in silico more rapidly and cheaply than could be achieved in a wet lab. Recent advancements in single-cell RNA sequencing (scRNA-seq) allow the capture of high-resolution snapshots of cell states as they transition along temporal trajectories. Using these high-throughput datasets, we can train computational models to generate in silico ‘synthetic’ cells that faithfully mimic the temporal trajectories. Here we present OneSC, a platform that can simulate synthetic cells across developmental trajectories using systems of stochastic differential equations govern by a core transcription factors (TFs) regulatory network. Different from the current network inference methods, OneSC prioritizes on generating Boolean network that produces faithful cell state transitions and steady cell states that mimic real biological systems. Applying OneSC to real data, we inferred a core TF network using a mouse myeloid progenitor scRNA-seq dataset and showed that the dynamical simulations of that network generate synthetic single-cell expression profiles that faithfully recapitulate the four myeloid differentiation trajectories going into differentiated cell states (erythrocytes, megakaryocytes, granulocytes and monocytes). Finally, through the in-silico perturbations of the mouse myeloid progenitor core network, we showed that OneSC can accurately predict cell fate decision biases of TF perturbations that closely match with previous experimental observations.

## Introduction

Cell fate decisions are governed, in part, by the set of interactions between transcription factors (TFs) and their target genes, which collectively form gene regulatory networks (GRNs) (Karlebach and Shamir, 2008; Cahan *et al*., 2021; Almeida *et al*., 2021; Churko *et al*., 2018; Moignard *et al*., 2015). Inferring GRNs and using them to model cell behavior is widespread in development biology (E. Y. Su *et al*., 2022; Alanis-Lobato *et al*., 2024), cancer biology (Seçilmiş *et al*., 2020; Keyl *et al*., 2023), drug development (Alvarez *et al*., 2018) and cell fate engineering (Cahan *et al*., 2014; Hartmann *et al*., 2018; Rackham *et al*., 2016). There are three commonly used yet distinct approaches to reconstructing GRNs. In one approach, the set of genes bound and potentially regulated by a TF, its regulon, is determined with methods such as ChIP-seq (Park, 2009) and CUT&Tag (Kaya-Okur *et al*., 2019). This approach is feasible when studying biological systems comprised of relatively few TFs for which antibodies are available but is infeasible otherwise. A second approach infers GRNs by mapping TF binding site (TFBS) motifs to accessible chromatin (Buenrostro *et al*., 2013; Song and Crawford, 2010) in cis regulatory elements of putative target genes (Pranzatelli *et al*., 2018; Yan *et al*., 2020). A third approach is to infer GRNs genome-wide based on the assumption that statistical association between TF and putative target gene expression implies a regulatory interaction, which can be achieved using bulk (Marbach *et al*., 2012) and single-cell RNA-seq data (Nguyen *et al*., 2021; Pratapa *et al*., 2020). The relative merits of these methodologies and efforts to integrate them have been reviewed elsewhere (Badia-I-Mompel *et al*., 2023; Kim *et al*., 2023).

In this study, we focus specifically on inferring GRNs from scRNA-seq in such a way to maximize the fidelity of GRN-generated expression states in comparison to real transcriptional states. Currently there are many methods to infer GRNs from single-cell expression profiles.

These methods can largely be grouped into several broad categories based on the main algorithms: tree-based methods (GENIE3 (Huynh-Thu *et al*., 2010), GRNBoost2 (Moerman *et al*., 2019)), correlation or information theory based methods (PPCOR (Kim, 2015), Epoch (E. Y. Su *et al*., 2022), LEAP (Specht and Li, 2017), PIDC (Chan *et al*., 2017), SCRIBE (Qiu *et al*., 2020)), regression based methods (SINCERITIES (Papili Gao *et al*., 2018), SINGE (Deshpande *et al*., 2022)), differential equations based methods (SCODE (Matsumoto *et al*., 2017), GRISLI (Aubin-Frankowski and Vert, 2020)) and Bayesian network based method (GRNVBEM (Sanchez-Castillo *et al*., 2018)). GRNs are typically evaluated based on regulatory edge recovery (precision and recall) rather than the extent to which inferred GRNs are able to operate as a functional dynamical system (i.e. can the GRN model or simulate certain biological phenomenal such as cell differentiation?). Here, we define functional GRNs as those with the following two properties: 1) capable of generating rich dynamical behaviors that reflect biologically relevant steady states (Ye *et al*., 2019; Guantes and Poyatos, 2008; Huang *et al*., 2022; Heydari *et al*., 2021; K. Su *et al*., 2022) and 2) capable of generating perturbation predictions (Heydari *et al*., 2021; K. Su *et al*., 2022).

Boolean networks can satisfy these conditions, and have successfully been used to model a wide range of biological phenomenon such as embryonic stem cell self-renewal (Dunn *et al*., 2019), transitions between pluripotent states (Yachie-Kinoshita *et al*., 2018), T-cell development (Heydari *et al*., 2021; Ildefonso and Finley, 2023), and *Drosophila melanogaster* segment polarity (Parmer *et al*., 2022). Previously RE:IN was developed to synthesize Abstract Boolean Networks (a set of concrete Boolean networks) that are consistent with experimental constraints and was used to identify the transcription factors circuit for modelling naïve pluripotency (Dunn *et al*., 2014). More recently, IQCELL used satisfiability modulo theory engine (Z3) (de Moura and Bjørner, 2008), a computational method that was also used in RE:IN (Dunn *et al*., 2014) and Hamey et al (Hamey *et al*., 2017), to identify Boolean logic functions that are constraint by the pseudo-time dynamics of binarized gene expressions (Heydari *et al*., 2021). The authors have demonstrated IQCELL’s capabilities of constructing Boolean networks that model early mouse

T-cell and red blood cells development. Although simple, Boolean networks are an attractive method to study functional gene regulation because they capture the essential sigmoidal step function for characterizing regulatory processes concentration levels (Schwab *et al*., 2020) even in the absence of difficult-to-determine kinetic parameters (Barbuti *et al*., 2020). Simulating changes in Boolean network states is computationally straightforward (Wang *et al*., 2012; Schwab *et al*., 2020) and thus facilitates performing dynamical simulations that produce testable predictions (Wang *et al*., 2012; Schwab *et al*., 2020). However, generating a functional Boolean network that produces attractor states mimicking the real data is not a trivial feat. From a previous benchmark of a dozen GRN inference methods using single-cell expression profiles (Pratapa *et al*., 2020), it showed that many of the methods, while not designed specifically for inferring functional Boolean networks, cannot produce Boolean networks with high simulation fidelity that exhibit the same number of steady states identical to those of the original models.

Here we present OneSC (**<u>One</u>** tool to **<u>S</u>**imulate **<u>C</u>**ells), a computational platform to simulate cell state transitions observed in single-cell expression data using a system of stochastic differential equations guided by an inferred functional GRN. BoolODE (Pratapa *et al*., 2020) has previously demonstrated the feasibility of simulating realistic synthetic single-cell expression profiles across developmental trajectories using a GRN and a system of stochastic differential equations that fundamentally represent a set of pre-defined Boolean logic for transcription regulations.

Extending the idea of simulating synthetic single-cell expression profiles using a functional gene regulatory network, OneSC has two main components 1) a more scalable computational method to simulate single-cell expression profiles similar to that of BoolODE and 2) a computational method to infer a GRN that maximizes the fidelity of GRN-generated expression states in comparison to real transcriptional states. Similar to BoolODE, OneSC also uses stochastic differential equations to simulate expressions of genes governed by transcription regulations from the network and a set of pre-defined Boolean logic. Improving upon BoolODE, OneSC’s simplified stochastic differential equations to model regulation allows OneSC to be more scalable and faster when simulating dense networks, and OneSC allows for native perturbation functionality.

In the other part of OneSC, we focus specifically on inferring GRNs from scRNA-seq data in such a way to maximize the fidelity of GRN-generated expression states in comparison to real transcriptional states using the OneSC simulator. Previously, a landmark benchmark study (using BoolODE and the BEELINE platform) reported that networks inferred from state-of-the art inference methods suffered from low simulation fidelity and were unable to generate the same number of steady states as found in the ground truth networks (Pratapa *et al*., 2020). To address this, our primary goal with OneSC was to improve upon the low simulation fidelity issues of current GRN inference methods. Our approach was to use additional information from standard scRNA-seq processing results (Luecken and Theis, 2019) such as cell clusters and pseudotime annotations, in addition to gene-level expression estimates, to construct Boolean networks that capture the cell states transitions trajectories and generate attractor states that match the terminal cell types observed in the single-cell dataset.

OneSC has two notable features that distinguish it from the recently published IQCELL method (Heydari *et al*., 2021) . First, GRN inference can be performed on multiple trajectories such as bifurcation or trifurcation without needing to infer the trajectory specific GRNs. Second, OneSC simulation platform allows for more continuous simulation of gene expressions than discrete asynchronous Boolean update.

In this work, we benchmarked OneSC’s GRN inference performance with over a dozen of current GRN inference methods using synthetic data on BEELINE platform (Pratapa *et al*., 2020), finding that OneSC networks have higher F1 scores. More importantly, we find that simulations performed with OneSC-generated networks achieve a higher similarity to real data.

We also find that OneSC’s stochastic differential equation simulation platform, though very similar to BoolODE, is more computationally scalable. To illustrate how OneSC could be used to derive biological insights, we applied it to real single-cell expression data of mouse myeloid progenitors cells (Paul *et al*., 2015). This analysis produced functional Boolean networks that recaptured normal myelopoietic differentiation trajectories, and in silico perturbations resulted in cell state compositions that largely match in vivo and in vitro genetic perturbations. In conclusion, we show that OneSC, coupling GRN inference and network simulation functionalities, is a useful tool to model differentiation and predict consequences of TF perturbations.

## Methods

### An overview of using OneSC to infer GRNs and to simulate transcriptional trajectories

The first major function of OneSC is to infer GRNs from scRNA-seq data. OneSC requires three minimal inputs to perform this task: normalized single-cell expression profiles, cluster/cell type annotation, and pseudotime assignments (**Fig 1A**). Given these, OneSC first constructs a cluster/cell type transition graph that reflects the relationship between clusters based on pseudotemporal order and expression profile similarity (**Fig 1B, Supplementary Info S1A**).

**Figure 1.**
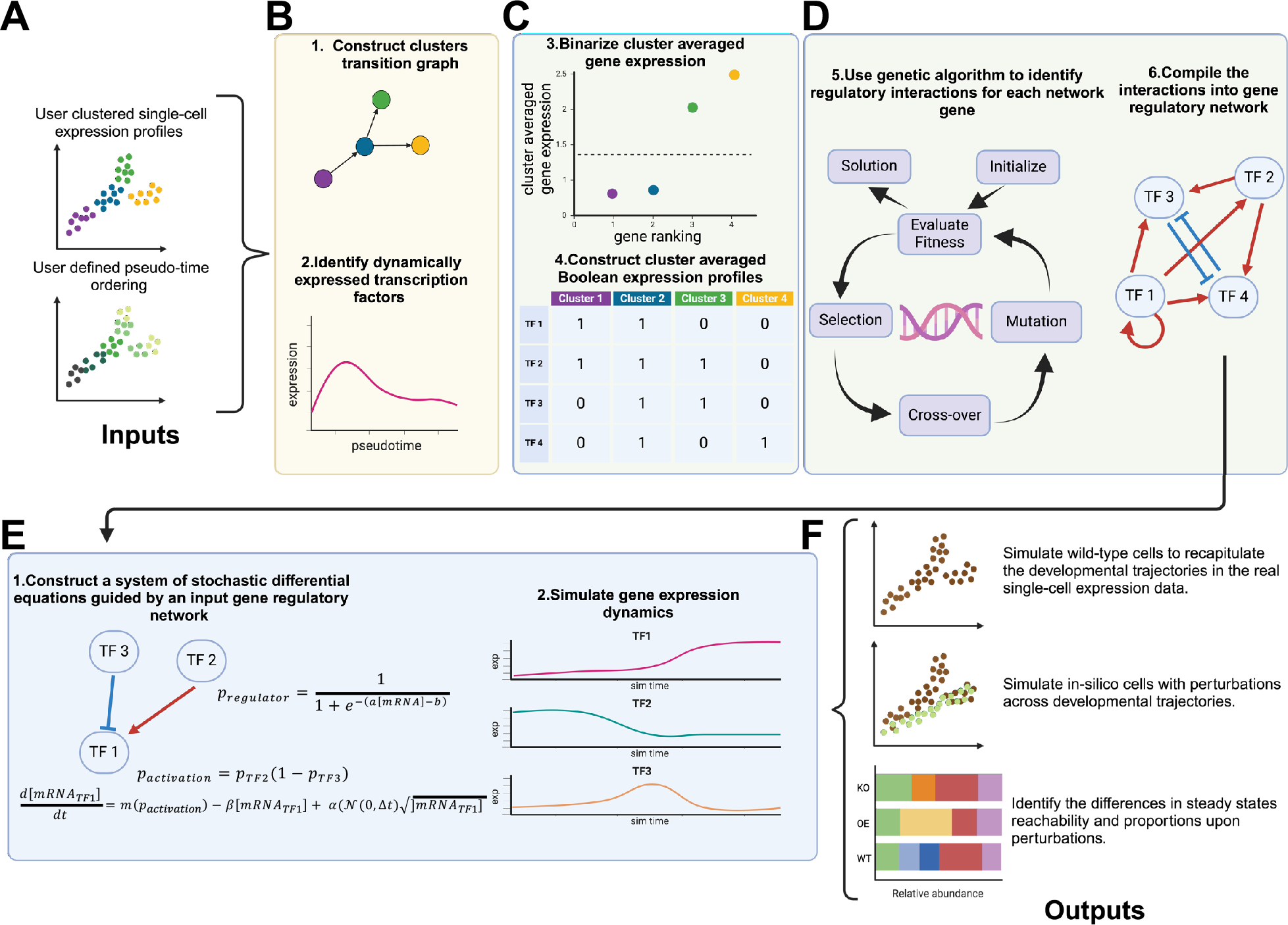
An overview of OneSC workflow. (A) Using the single-cell expression profiles and the inputted cell type clusters annotations with pseudotime information, (B) OneSC first constructs the cluster transition graph summarizing the cell state transitions observed in the single-cell data. Users can also manually create the clusters transition graph without using OneSC’s built- in function. From the individual trajectories in the cell state transition graph, OneSC next identifies dynamically expressed transcription factors for the construction of the gene regulatory network. Users can also manually select a set of transcription factors. (C) Then, OneSC averages the gene expression profiles for each cell type cluster and binarized them into activity status (1 or 0 representing on or off). (D) For each gene in the network, OneSC uses genetic algorithm to identify a set of regulatory interactions between target gene and regulators such that the agreement between the activity status of the target gene and the simulated activity status of the target gene across cell states is maximized. Lastly OneSC compiles all the subnetworks for individual genes into a large network. (E) Using the inferred network, OneSC constructs a system of stochastic differential equations to simulate the gene dynamics. (F) Coupling the network inference and simulation, OneSC can generate simulated cells during normal cell state transitions or under perturbation to predict the shift in steady state reachability and proportion.

Second, OneSC identifies a core set of dynamically expressed TFs for each trajectory with a generalized additive model (E. Y. Su *et al*., 2022) (**Fig 1B, Supplementary Info S1B**). It is important to note that users can also construct cell state transition graphs and select important TFs manually, independent of OneSC’s helper functions or they can modify upon the results obtained from OneSC’s helper functions. Third, OneSC averages the expression of dynamic TFs in each cell type cluster and binarizes them into activity status: 1 or 0 (on or off) resulting in Boolean expression profiles (**Fig 1C, Supplementary Info S1C**). Fourth, OneSC uses genetic algorithm (GA) (Mirjalili, 2019), an evolutionary inspired metaheuristic optimization process that has previously been used in GRN inference (Barman and Kwon, 2018; Zhang *et al*., 2021; Trinh and Kwon, 2021; Chen *et al*., 2015) and refinement (Park *et al*., 2023), to identify regulatory interactions (i.e. activation, repression, or no regulation) between the potential regulators for each target gene (**Fig 1D, Supplementary Info S1D**). The goal of GA here is to identify a subnetwork of direct regulators that maximizes the agreement between simulated activity status of the target gene (driven by the subnetwork configuration and activity status of the direct regulators) and the activity status observed in the real data. Lastly, OneSC compiles all the inferred subnetworks for each target gene into one large functional GRN (**Fig 1D**).

The second major function of OneSC is to simulate transcriptional states from the inferred GRN and an initial state activity status profile. Alternatively, OneSC offers the flexibility for users to use their own curated gene regulatory networks and simulate them if the GRN conforms to the constraint that all nodes in the network have at least one regulator (including self-regulator).

OneSC achieves this with systems of stochastic differential equations that model the expression dynamics of each gene (**Fig 1E, Supplementary Info S2**). The expression value of target genes is regulated by the activity status of upstream TFs through a set of Boolean algebra (Shannon, 1940). With the use of Boolean algebra and differential equations, OneSC dynamically simulates continuous expression values between simple binary values (on and off) to overcome the limitation of binary output of asynchronous or synchronous Boolean updates (Barbuti *et al*., 2020). OneSC then generates synthetic expression profiles across simulation time that mimic pseudotemporal trajectories (**Fig 1F**). Furthermore, OneSC can natively perform in silico overexpression and knockout to explore how these perturbations impact trajectories, the reachability of terminal states, and cell type/cluster compositions (**Fig 1F**).

## Results

### OneSC accurately simulates expression trajectories

Simulating accurate expression trajectories is dependent on the quality of the GRN that is used. Therefore, we first assessed OneSC’s ability to infer GRNs using gold standard synthetic data and ground truth GRNs from the BEELINE platform (Pratapa *et al*., 2020) (**Supplementary Info S3**). We downloaded 10 sets of synthetic datasets (2000 cells, no drop-out) with gold standard GRNs, manually clustered the datasets using standard Scanpy (Wolf *et al*., 2018) pipeline, and use the provided Slingshot-computed pseudotime (Pratapa *et al*., 2020), which we provided as input to OneSC’s GRN inference function (**Fig S1-3**). Among the ten gold standard GRNs, there are six synthetic networks: dyn-BF (12 edges and 5 nodes), dyn-BFC (18 edges and 9 nodes), dyn-CY (6 edges and 5 nodes), dyn-LI (8 edges and 7 nodes), dyn-LL (19 edges and 18 nodes), dyn-TF (20 edges and 7 nodes) and three literature curated networks: GSD (79 edges and 18 nodes), HSC (30 edges and 11 nodes), mCAD (14 edges and 5 nodes) and VSC (15 edges and 8 nodes). Then we computed precision and recall based on comparison of inferred GRNs to the ground truth GRNs (**Fig S4A-B**) against 13 other GRN inference methods (GENIE3 (Huynh-Thu *et al*., 2010), GRNBoost2 (Moerman *et al*., 2019), PPCOR (Kim, 2015), PyEpoch (E. Y. Su *et al*., 2022), LEAP (Specht and Li, 2017), PIDC (Chan *et al*., 2017), SCRIBE (Qiu *et al*., 2020), SINCERITIES (Papili Gao *et al*., 2018), SINGE (Deshpande *et al*., 2022), SCODE (Matsumoto *et al*., 2017), GRISLI (Aubin-Frankowski and Vert, 2020), GRNVBEM (Sanchez-Castillo *et al*., 2018), IQCELL (Heydari *et al*., 2021)). To be as fair as possible in this comparison, we selected edge weight thresholds that optimized F1 for those methods that produce edge weights (all methods except OneSC and IQCELL).The mean F1 score, which is the harmonic mean of precision and recall, of OneSC’s GRNs was 0.61 which was as good as or higher than other GRN inference methods applied to the same data (**Fig 2A**). This result suggests that OneSC, without needing users to define thresholds post GRN inference, is comparable to other GRN inference methods when assessed on traditional GRN performance metrics (see also **Fig S4A- B**). In terms of runtime, OneSC is on average ranked as the second slowest method behind IQCELL (**Fig S4C**).

**Figure 2.**
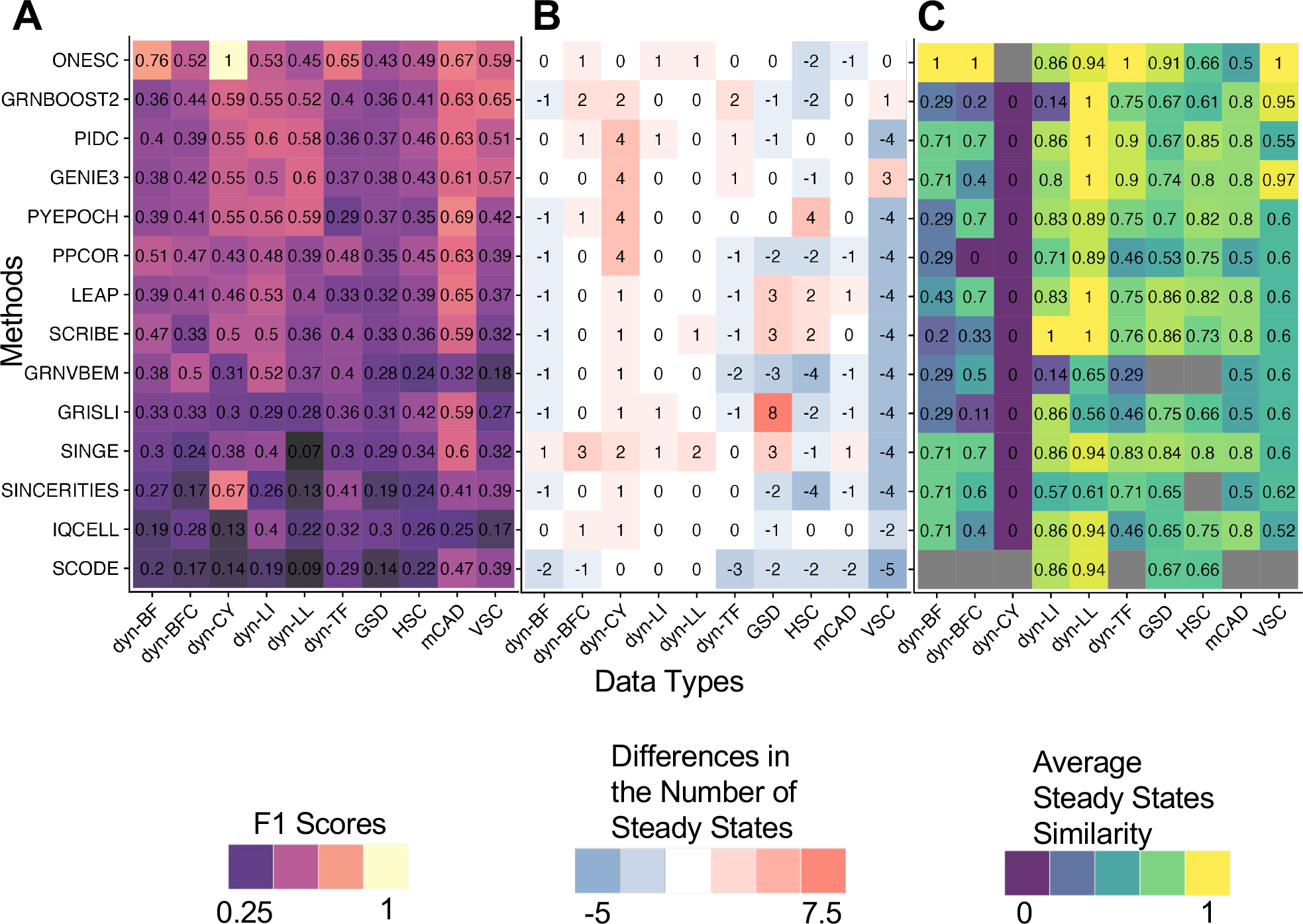
Benchmarking OneSC gene regulatory network inference with other methods using BEELINE platform. (A) Heatmap showing the maximum F1 scores of various GRN inference methods tested on 10 synthetic datasets with associating gold standard networks. (B) Heatmap showing the differences in the number of steady states generated by the inferred GRNs and gold standard networks using asynchronous Boolean updates. (C) Heatmap of the average similarity between Boolean activity profiles of the steady states generated from inferred networks and those from gold standard networks using asynchronous Boolean updates. Similarity score is calculated as the percentage of genes that match activity status between steady states generated using inferred network and those generated using gold standard network. Similarity score of one indicates perfect match between simulated profile from inferred network and gold standard steady state. For each gold standard network, the mean of the maximum similarities between gold standard steady states and inferred network’s steady states are used to represent the overall similarity between inferred GRN’s steady states and gold standard network’s steady states. In the dyn-CY dataset the average steady state similarity score is assigned a zero if the inferred network has a steady state because dyn-CY gold standard network does not have a steady state. The grey color box indicates that no steady state was found in the inferred network.

Next, we assessed the robustness of OneSC to parameter values (**Supplementary Info S4**). First, we explored how cluster assignment impacts GRN inference. We performed Leiden clustering on the benchmark data by finding the highest clustering resolution between 0.1 and 0.75 (0.05 increment) that still ensures there is at least one gene with substantial mean expression difference (>1.5) between all cluster pairs. This ensures that each cluster has a unique Boolean activity profile. To test the stability of the inference performance with when there are over-clustering or under-clustering, we randomly split or merged clusters, performed network inference, and then determined the precision and recall. From this analysis, we found that there can be as large as a 50% decrease in F1 score when deviating from the optimal cluster assignment (**Fig S5A**). Therefore, we advise users to run OneSC with cell clusters that are meaningful and with distinct expression profiles given the biological context. Next, we tested the impact of varying the ideal number of edges and the number of GA generations but did not see noticeable differences in the performance if we adjusted other user tunable parameters such as the ideal number of network edges (**Fig S5B**) or the number of generations for the genetic algorithm optimization which is linearly correlated with runtime (**Fig S6A-B**).

Having determined the stability and performance of OneSC’s GRN inference approach, we next assessed the extent to which OneSC-inferred GRNs recapitulate the same number of steady states as in the gold standard data following the assessment used by the BEELINE benchmarking study (Pratapa *et al*., 2020). To perform this analysis, we ran 5, 000 repressor centric Boolean asynchronous simulations for each inferred GRN (given the same initial state that was used in BoolODE to generate the synthetic data) and then determined the number of distinct steady states reached. We found that OneSC generated the correct number of steady states in 5 out of 10 gold standard datasets with dyn-BFC, dyn-LL, dyn-LI networks generating one more steady state than gold standard networks, and HSC, mCAD networks generating two and one fewer steady states than gold standard networks respectively (**Fig 2B**). Next, we compared the similarity of inferred networks’ steady states to the gold standard steady states by calculating and finding the maximum percent agreement between the Boolean expression profiles from inferred networks’ steady states and those of the gold standard. This process was performed for each of the gold standard steady state in the synthetic dataset and the average percent similarity of all the steady states in the synthetic dataset was calculated to show the overall similarity in terms of steady state reachability between inferred networks and gold standard networks (**Fig 2C**). We found that OneSC networks generated steady states were identical to the steady states of gold standard networks in dyn-BF (2 steady states), dyn-BFC (1 steady state), dyn-TF (3 steady states), dyn-CY (0 steady state) and VSC (5 steady states) structures (**Fig 2C**). OneSC networks also generated steady states that were highly similar (average percentage of gene activity status agreement >= 0.86) to gold standard networks’ steady states in dyn-LI (1 steady state) and dyn-LL (1 steady state), GSD (2 steady states) synthetic cell structures. We found overall lower average steady states similarity in OneSC’s HSC (4 steady states) and mCAD (2 steady states) inferred networks (**Fig 2C**) likely due to the fact that these networks generated fewer steady states than gold standard networks (**Fig 2B**).

Overall, we found that networks generated using OneSC generally produced networks with the highest average steady states similarity (mean 0.87) and second lowest divergent number of steady states (absolute mean 0.6) right underneath IQCELL (absolute mean 0.5). Taken together, this evaluation supports the notion that OneSC can capture enough of the key regulatory structure of the underlying GRNs to preserve dynamical properties such as the reachability and similarity of steady states.

### OneSC simulations scale linearly with number of nodes in the network

The second central aspect of OneSC is the capability to generate single cell expression states that reflect pseudotemporal trajectories using a Boolean network backbone. OneSC’s simulation platform is heavily inspired by a current method, BoolODE (Pratapa *et al*., 2020). Currently, BoolODE has two areas for improvement. The first is native support for simulating expression states upon network perturbation (i.e. knocking out or over-expressing genes). To address this, we have designed OneSC such that it is straight-forward for users to test the impact of perturbing on pseudotemporal trajectories (see next section for example applications).

The second missing feature is computational scalability with the number of regulators per target gene. For example, it is infeasible to simulate expression trajectories with BoolODE for networks in which there are ten or more regulators per target gene because the time complexity of the simulation grows exponentially with respect to the number of regulators (Dibaeinia and Sinha, 2020). To reduce computational workload, we first eliminated the differential equations to model protein concentration dynamics. Second, we simplified regulatory functions using Boolean algebra such that the time complexity with respect to the number of regulators is linear.

We benchmarked OneSC simulator with BoolODE and BoolODE soft-heaviside (a faster version of BoolODE that also uses Boolean algebra) using 20 different unique network types across different network sizes {5, 10, 15, 20} and network densities {0.2, 0.4, 0.6, 0.8, 1} (**Fig 3A-B, Supplementary Info S5**). There are ten random different network structures for each unique network configuration. For each random network, we ran five simulations with 5000 simulation steps turning off multi-core parallel processing (**Fig 3A**). With smaller networks (i.e. those with 5 and 10 nodes and fewer than 75 edges), simulations with BoolODE and BoolODE soft-heaviside complete more rapidly than OneSC (**Fig 3Ai, ii**). However, at higher network size and density (i.e. those with 15 and 20 nodes and with more than 75 edges), OneSC simulations complete faster than those of BoolODE and BoolODE soft-heaviside (**Fig 3Aiii, iv**). We found that BoolODE and BoolODE soft-heaviside failed to simulate at certain edge counts (120 edges with 15 nodes, 140 edges with 20 nodes) (**Fig 3Aiii, iv**) as previously reported (Dibaeinia and Sinha, 2020). Another observation to note is that while BoolODE’s runtime is sensitive to the number of edges and the number of nodes in the network, OneSC’s runtime is primarily correlated with the number of nodes and is less sensitive to the number of edges in the network. When using multiple core parallelization to perform simulations, BoolODE and BoolODE soft-heaviside improved in terms of runtime for smaller networks: 5 node networks (**Fig 3Bi**), 10 node networks (**Fig 3Bii**), 15 nodes networks with less than 120 edges (**Fig 3Biii**), 20 nodes network with less than 140 edges (**Fig 3Biv**) while the runtime remained the same as single-core usage for OneSC. Consistent with single-core results, BoolODE and BoolODE soft-heaviside fail to simulate larger and denser networks (15 nodes network with more than 120 edges, 20 nodes network with more than 140 edges) (**Fig 3Biii, iv**).

**Figure 3.**
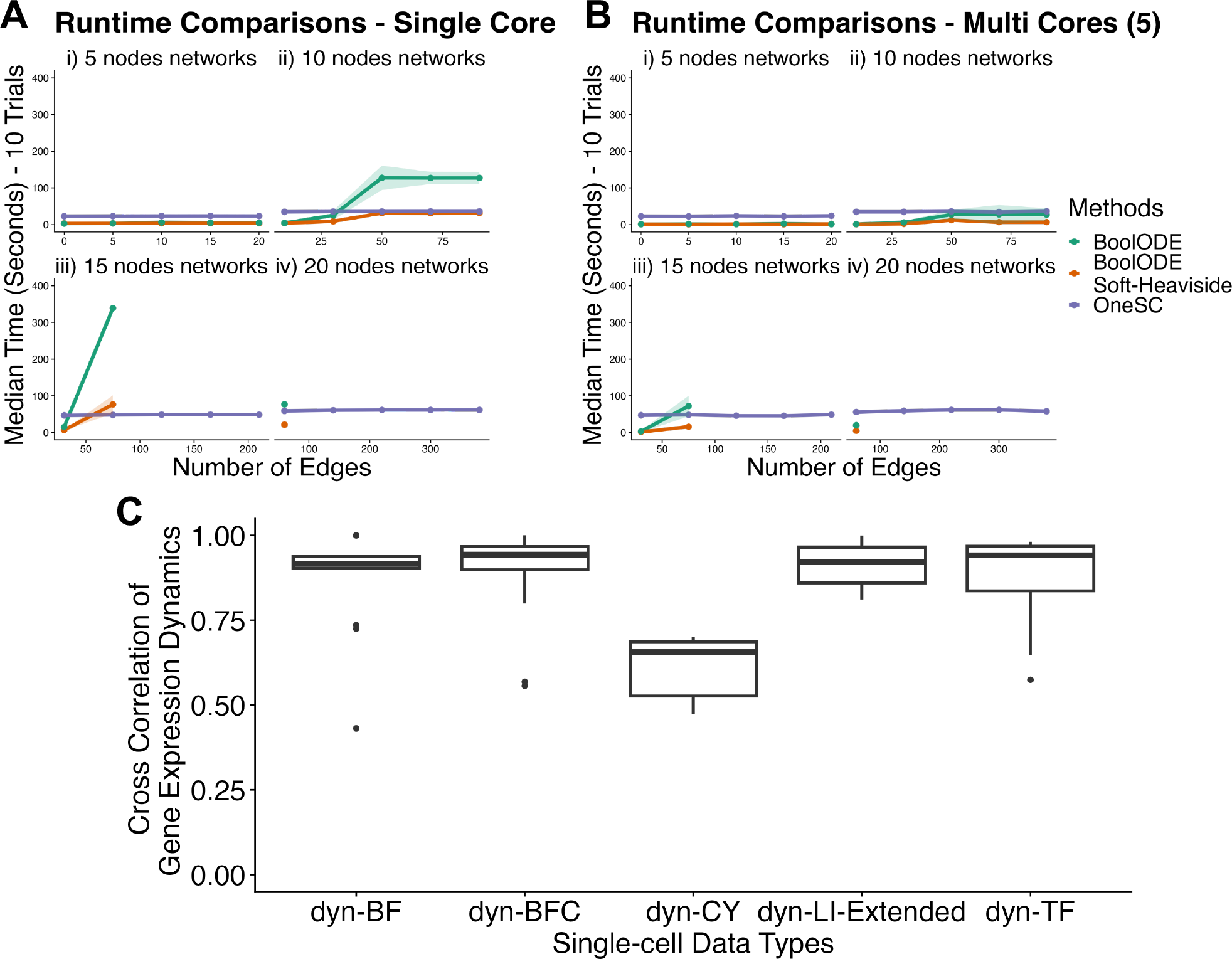
Runtime and similarity comparisons between OneSC simulator and BoolODE. (A) The median runtime for OneSC, BoolODE and BoolODE (soft-heaviside) to simulate 10 random networks generated across different network sizes (i) 5, (ii) 10, (iii) 15, (iv) 20 and different network densities (0.2, 0.4, 0.6, 0.8, 1) without using parallelization. The shaded area represents the range of runtime across 10 random networks simulations. If there is no data point for a particular network configuration, it means the simulation was not able to run. (B) The median runtime for OneSC, BoolODE and BoolODE (soft-heaviside) to simulate 10 random networks generated across different network sizes (i) 5, (ii) 10, (iii) 15, (iv) 20 and different network densities (0.2, 0.4, 0.6, 0.8, 1) with parallelization (5 cores). (C) Cross-correlation of network gene expression dynamics between simulated data from OneSC and BoolODE. If the dataset has more than one trajectory, the cross-correlations of genes expressions are computed individually for each trajectory. To ensure the simulated data from BoolODE reached steady state, we re-simulated dyn-LI gold standard network using BoolODE and extended the simulation time from 5 to 7.

Next, we sought to determine the extent to which our reduction in the complexity of simulation equations and removal of protein simulation equations impacted the fidelity of OneSC’s simulations. To assess this, we compared OneSC’s simulated expression trajectories based on the gold standard GRNs to those generated by BoolODE, finding a high degree of cross-correlation between them (**Fig 3C**). The dyn-CY (cyclic dynamical structure) simulation was the only exception and was likely due to differences in the periodicity of cyclic gene expression.

Furthermore, all the distinct clusters in the single-cell data generated by BoolODE are also observed from the single-cell data generated by OneSC (**Fig S7-8**). Taken together, these results show that using one OneSC for expression state trajectory simulations is more computationally tractable than other, similar tools such as BoolODE.

### Predicting the impact of TF perturbation on myelopoiesis

To demonstrate the utility of OneSC on real single-cell expression data, we applied it to a scRNA-seq data set of mouse myeloid progenitors (Paul *et al*., 2015) to infer a core TF GRN circuit, to simulate differentiation trajectories, and to explore the consequences on myelopoiesis of perturbing these TFs. Prior to running OneSC, we clustered the cells and annotated as: common myeloid progenitor (CMP), erythrocytes, megakaryocytes, monocytes, granulocytes, granulocyte monocyte progenitor (GMP), megakaryocyte erythrocyte progenitor (MEP) based on marker gene expression (**Supplementary Table 1**) (**Fig S9A**). Then, we computed pseudotime using diffusion pseudotime (Haghverdi *et al*., 2016) and manually set the CMP cluster as the start of the trajectory and erythrocytes, megakaryocytes, monocytes and granulocytes as the terminal cell states to construct the cell state transition graph (**Fig 4A**). In this dataset, megakaryocytes have an earlier pseudotime than their nominal progenitor MEP (**Fig S9A, B**) consistent with recent recognition of a direct differentiation from CMP to megakaryocytes that bypasses an MEP intermediate stage in some contexts (Miyawaki *et al*., 2017; Sanjuan-Pla *et al*., 2013; Notta *et al*., 2016). Next, we used OneSC’s function to identify dynamically expressed TFs, which yielded 12 genes, all of which have been implicated in various aspects of hematopoiesis (**Fig 4B, S9C**). Finally, we used OneSC to infer a GRN (**Fig 4C**), which we then used to simulate 200 simulation runs (i.e. synthetic single cell trajectories) with 1800 simulation steps (analogous to pseudotime ordering), recapitulating the native cell state trajectories (**Fig 4D, 4E**). The initial state was defined based on the activity status profile of the CMP cluster. Most of the 200 simulation runs stabilized into four cell states (i.e. Boolean activity status profiles) that are identical to monocytes, granulocytes, megakaryocytes (MK) or erythrocytes (**Fig 4D, S10**). Intermediate cell states such as MEP and GMP are occupied for brief intervals in some simulation runs, suggesting that the simulations follow developmental trajectories according to those in the training data (**Fig S10**). Looking at selected TFs expressions between simulated and real single-cell expression data (**Fig S11, S12**), we saw a high degree of similarity between the two. For instance, both simulated cells and real monocytes cells express *Irf8*, an important transcription factor for murine monocyte differentiation (Kurotaki *et al*., 2013), while granulocytic cells express *Cebpe*, a regulatory of granulopoiesis

**Figure 4.**
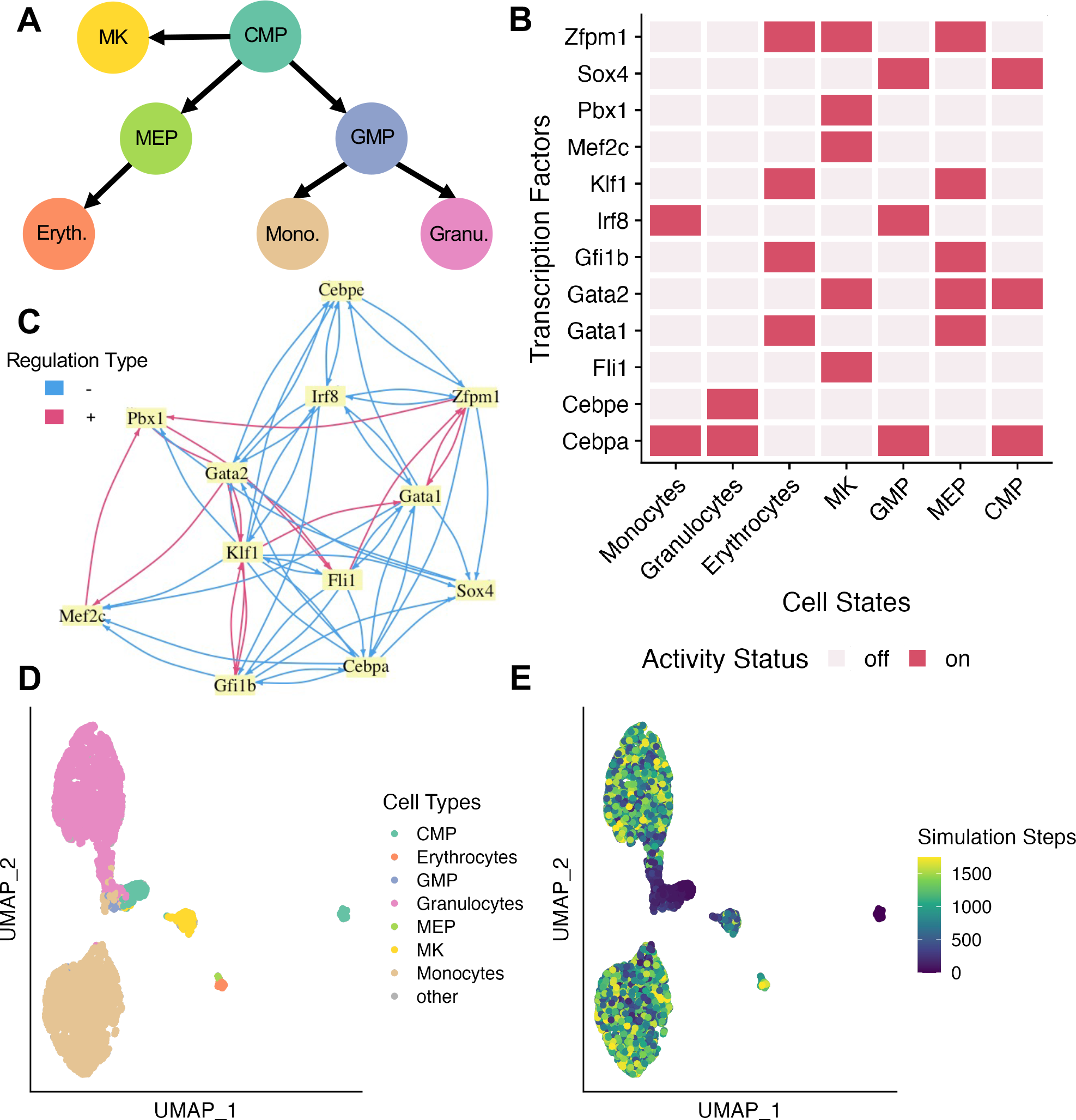
Application of OneSC to model developmental trajectories in mouse myeloid progenitor cells. (A) The cell state transition graph for mouse myeloid progenitor cells. (B) Heatmap representing the Boolean activity profiles of the 12 dynamically expressed transcription factors across 7 cell states defined in the dataset. (C) OneSC’s inferred transcription factor circuit from mouse myeloid progenitor cells. Blue edges represent repression and red edges represent activation. (D) UMAP embeddings of synthetic cells across 200 simulation runs (sampled at every 50 time-steps). The cells are labelled by the cell type with the lowest distance in Boolean activity profiles. Trajectories labelled as “other” are simulation runs in which the steady state Boolean activity profiles do not match perfectly with any of the Boolean profiles of the terminal cell types (MK, Erythrocytes, Granulocytes, Monocytes). (E) UMAP embeddings of the synthetic cells across 200 simulation runs (sampled at every 50 time steps) colored by the simulation time steps ordering. The black color represents the earliest time point in the simulation and the yellow color represents the latest time point sampled form the simulation.

(Lekstrom-Himes, 2001) (**Fig 4D, S9A, S11, S12**). On the other trajectory, both simulated and real erythrocytes express *Klf1*, an essential regulator for erythropoiesis (Siatecka and Bieker, 2011) and megakaryocytes express *Pbx1* (**Fig 4D, S9A, S11, S12**).

Next, we explored the extent to which simulations with GRNs inferred by other methods mimicked myelopoiesis trajectories using OneSC’s simulator. Many of the GRN inference methods output an edge-weighted network instead of a concrete Boolean network. To convert those networks into concrete Boolean networks, we first identified the thresholds at which the resulting Boolean network would have the same density as OneSC’s GRN or less if it exceeds the maximum possible density. We found that OneSC network is the only network capable of reaching all the steady states using the Boolean simulation (**Fig 5A**). While not surprising given that most GRN inference methods were not designed or optimized for this task, our results show that OneSC fills the need for inference of executable Boolean networks with dynamical properties that faithfully mimic cell type transitions in single-cell expression data.

**Figure 5.**
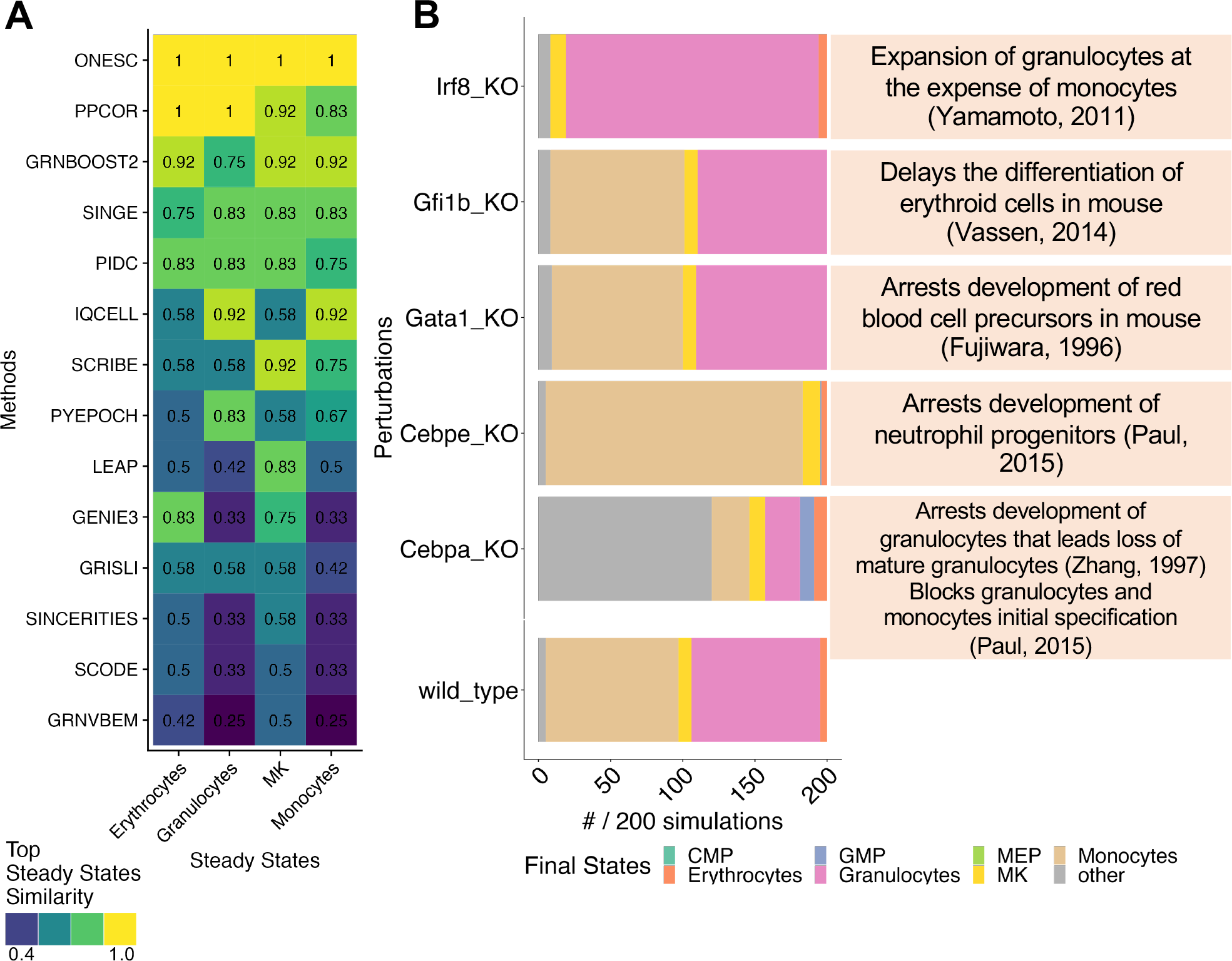
Assessment of OneSC network’s simulation fidelity and perturbation predictions. (A) Heatmap comparing the steady state similarities of the networks generated from OneSC and other gene regulatory network inference methods for Monocytes, MK, Granulocytes and Erythrocytes. Similarity for each cell type is measured by highest percent agreement between Boolean activity profiles of the simulated steady states and those from the real data. (B) Barplot showing the proportion of steady states from OneSC’s knockout simulations of *Irf8, Gfi1b, Gata1, Cebpe, Cebpa* and no perturbation simulation (wild_type) (left). The qualitative observations from previous published in-vivo knockout experiments for *Irf8, Gfi1b, Gata1, Cebpe, Cebpa* (right).

Several of the TFs in our myelopoiesis GRN have been previously explored by generating transgenic mouse knockouts or by altering their expression. Therefore, we next explored how the biological consequences of these experimental perturbations related to perturbations that we could perform *in silico*. We simulated cellular trajectories upon knocking out and overexpressing each TF and recorded the final state composition (**Fig 5B, S13A-B**). We examined five TFs for which we could find literature in which the TF was knocked out or perturbed in a relevant system.

First, we looked at the simulation of *Irf8* knockout. We found that there was an increase in granulocyte and decrease in monocyte final states (**Fig 5B**). These results are largely consistent with experimental data. Mice lacking the *Irf8* gene (*Irf8*^-/-^ mice) have disproportionate expansion of neutrophils at the expense of monocytes/macrophages (Yamamoto *et al*., 2011; Tsujimura *et al*., 2002; Scheller *et al*., 1999). In our simulation, *Irf8* overexpression influenced all the simulation runs to arrive at the monocyte final state (**Fig S13A**) suggesting that *Irf8* is a key driver in the monocytic development. When *Gfi1b* and *Gata1* were knocked out, there were fewer erythrocytes in the knockout simulations compared to that of wild-type simulation (**Fig 5B**). This is in agreement with prior experiments in which *Gfi1b* knockout delays erythroid differentiation (Vassen *et al*., 2014). Similarly, *Gata1* knockout arrests development of red blood cell precursors (Fujiwara *et al*., 1996).

Our simulations of *Cebpe* knockout mirrored prior experiments in which loss of *Cebpe* arrests neutrophil progenitor development (Paul *et al*., 2015) (**Fig 5B**). *Cebpa* knockout was previously found to arrest development of granulocytes and blocks granulocytes/monocytes initial specification (devoid of GMP) leading to a loss of monocytes and granulocytes (basophils and neutrophils) (Zhang *et al*., 1997; Paul *et al*., 2015). In-silico *Cebpa* knockout led to a drastic decrease of granulocyte and monocyte final state proportion, an increase in the number of GMPs and ’other’ cell types (i.e. cells that do not completely match the Boolean activity status profiles of any cell states) (**Fig 5B**).

We have also tested other in-silico perturbations without prior experimental verifications. For instance, when *Klf1*, an important transcription factor regulating the erythropoiesis (Tallack *et al*., 2010; Orkin and Zon, 2008), was overexpressed in-silico, there was an increase of erythrocytes as final states (**Fig S13A**). On the other hand, when we knocked out *Klf1* in-silico, there was a loss of erythrocytes in the final states (**Fig S13B**). Next we investigated the effect of perturbing *Zfpm1* or *Fog1*, an required transcription factor for the development megakaryocytic and erythrocytic lineage (Mancini *et al*., 2012). When *Zfpm1* was overexpressed, there were more final states that resembled megakaryocytes or erythrocytes than wild-type simulation (**Fig S13A**). On the other hand, when *Zfpm1* was knocked out, there was a decrease of final states that resemble megakaryocytes or erythrocytes (**Fig S13B**). Taken together, these results show that OneSC can recapitulate the expression trajectories of mouse myeloid differentiation and can accurately predict the cell impact of genetic perturbations on steady state composition.

## Discussion

With the increase availability of scRNA-seq datasets of various biological systems, there has been a vast amount of interest in using these resources to infer the underlying transcriptional circuits and to develop a method to simulate the dynamics of these networks for predictions. Here we present OneSC, an open-source Python package that can infer executable GRNs to faithfully recapitulate the developmental trajectories of biological systems and can stochastically simulate synthetic single-cell expression profiles that resemble single cells from real datasets.

We first benchmarked OneSC’s GRN inference capability using synthetic gold standard dataset from BEELINE platform (Pratapa *et al*., 2020) against other published GRN inference methods. Our results reveal that given the right clustering, OneSC can infer GRNs with precision, recall and F1 scores that match the highest performing concrete networks achievable by existing GRN inference methods through F1 maximization thresholding scheme. Different from most of the current GRN methods, OneSC does not require users to select a threshold post GRN inference to form a concrete network. This is an important feature because there is often no gold standard network for the user to identify the thresholds that maximize the F1 score of the inferred networks. Moreover, among the plethora of GRN inference methods, OneSC stands out for its capabilities in creating concrete Boolean networks that recapitulate the steady states observed in the training single-cell data, indicating that OneSC networks retain the essential regulations governing developmental processes. Secondly, OneSC’s simulator, heavily inspired by existing single-cell simulator BoolODE (Pratapa *et al*., 2020), overcame the scalability issue of BoolODE (Dibaeinia and Sinha, 2020) via the simplification of transcription regulation functions and the removal of protein production equations. OneSC has shown to be able to simulate denser and larger networks that BoolODE cannot simulate. Despite the simplification of equations for transcriptional regulations, OneSC can still generate different complex dynamical single-cell data types such as bifurcation or trifurcation. OneSC simulation platform also offers the flexibility for users to input their own gene regulatory networks. Lastly, we applied OneSC platform to real mouse myeloid single-cell dataset (Paul *et al*., 2015) and have demonstrated its capability of generating an executable GRN that captures the cell state transitions of various developmental trajectories. We have also demonstrated the predictive capabilities of OneSC by performing in-silico perturbations of key transcription factors and saw the changes in final cell states proportion largely match with experimental results in literature.

IQCELL is another integrative platform that performs Boolean network inference on single-cell data and simulates network with or without in-silico perturbations (Heydari *et al*., 2021). IQCELL uses the Z3 engine to find the optimal logical rules, or regulatory functions, for each gene (Hamey *et al*., 2017) based on candidate interactions obtained through mutual information that follow experimental data when executed as logical gates. With the inferred GRN, IQCELL uses asynchronous Boolean simulator to simulate under normal or perturbed condition (Heydari *et al*., 2021). Despite some similarities, there are several key differences between IQCELL and OneSC. First, OneSC uses an optimization method, genetic algorithm, to identify the best regulatory rules for each gene such that when simulated, the agreement between simulated and real activity status of target genes are maximized at cell type/cluster resolution instead of at single-cell resolution. Inferring GRNs at cell type/cluster resolutions via averaging cluster cells’ expressions, although loses single cell resolution, alleviates the issue of high noise due to drop- outs in scRNA-seq (Kim *et al*., 2020; Kharchenko *et al*., 2014; Hwang *et al*., 2018; Murphy and Skene, 2022). Second, OneSC uses a continuous simulation system that allows users to simulate the continuous gene expression dynamics (including partial on and partial off) while IQCELL uses discrete asynchronous Boolean update that has two states (on and off). Lastly, IQCELL was only tested in linear developmental trajectories. The authors of IQCELL recommend users to infer a candidate GRN for each trajectory if there are multiple trajectories in the dataset (Heydari *et al*., 2021). This restriction limits the simulation and predictive capability of IQCELL when applied to single cell datasets with multiple trajectories and steady states. On the other hand, OneSC generates a Boolean network that encompasses multiple developmental trajectories allowing OneSC to simulate all the trajectories at once using stochastic differential equations and make perturbation predictions that model changes in cell fate decisions.

There are several limitations and caveats to our computational framework. First, as shown in our benchmarking results, the performance of OneSC network inference is heavily dependent on the clustering/cell typing provided by the user. To utilize OneSC to its maximum capabilities, we recommend that users cluster cells that reflect distinct cell pseudotemporal states with distinct Boolean activity profiles of the network genes. Second, while OneSC performs well at recovering structural edges from synthetic gold standard networks, OneSC is designed to infer regulatory interactions that are functional but not necessarily be direct (Dunn, 2019). Third, OneSC fundamentally infers Boolean networks to study cell state transitions during development and to predict in-silico perturbations. Despite being an interpretable qualitative model of gene regulatory interactions that can generate predictive hypothesis via dynamical simulations (Wang *et al*., 2012; Barbuti *et al*., 2020), Boolean networks are limited by their discretization bottleneck (only on or off status) (Delgado and Gómez-Vela, 2019). OneSC tries to mediate this issue via the adoption of BoolODE-like simulation scheme that allows partial activation or partial repression of regulators. However, we recognize that there are other ways to perform perturbation predictions focused on a gene regulatory network that are not Boolean centric. For instance, there are regression based methods such CellOracle (Kamimoto *et al*., 2023) and Dynamo (Qiu *et al*., 2022), and differential equation based methods such as SERGIO (Dibaeinia and Sinha, 2020), PeTTSy (Domijan *et al*., 2016), RACIPE (Huang *et al*., 2018) and Dictys (Wang *et al*., 2023). There are also non-network dependent generative modelling methods such PRECIENT (Yeo *et al*., 2021). Furthermore, with the increasing prevalence of perturb-seq data (Dixit *et al*., 2016), there are new machine learning and deep learning methods (Roohani *et al*., 2023; Lopez *et al*., 2022) for prediction of perturbations benchmarked by Kernfeld et al (Kernfeld *et al*., 2023). Fourth, the input cell state transition graph has to be hierarchal and non-cyclic with one initial state and at least one terminal state. The later cell states on the developmental hierarchy cannot have a transitional edge to earlier cell states on the developmental hierarchy. In the case of cyclic transition graphic structure, we recommend the users to break it into linear structure before using as input in OneSC. Fifth, although we have developed simple helper functions to automatically construct the cell state transition graph based on pseudotime ordering and cluster expression similarity, we recognize that these functions may not perform well for all biological systems. Therefore, when the outputted state transition graph is incorrect, users should manually create the cell state transition graph with the help from tools specialized for this task such as PAGA (Wolf *et al*., 2019) or CellRank (Lange *et al*., 2022). Sixth, steady state reachability depends on the set of TFs in the GRN, and therefore this set may need to be expanded when simulations do not reflect real steady states. Lastly, with the emergence of single-cell multi-omics (Dimitriu *et al*., 2022; Lee *et al*., 2020), we are capable of elucidating more than just transcriptomic signals. For future work, we plan on developing pipelines to identify likely or unlikely regulators a priori via the integration of multi-omics data and use those as constraints to fine-tune the inferred networks.

## Conclusion

In conclusion, we present a computational tool, OneSC, that performs two main tasks. First, it infers executable gene regulatory Boolean networks from single-cell expression profiles, inferred pseudotime and cluster/cell type annotation given by the users. Second, OneSC performs dynamical simulations that generate synthetic single cells to mimic trajectories and cell state transitions. Coupling these two main functions, users can perform *in silico* gene perturbations to predict shifts in terminal cell states, blockage or promotion of certain lineages and changes in gene expression dynamics. To promote accessibility of OneSC (https://github.com/CahanLab/oneSC), we have made the code freely available under an Open-Source license and as an easily installable Python package.

## Acknowledgements

We would like to thank members of the Cahan lab for providing feedback and support, and additionally to Eric Kernfeld and Alex Chen for helpful discussions on the manuscript. This work was funded by National Institutes of Health (NIH R35GM124725 and R01CA262802) and Maryland Stem Cell Research Foundation (2022-MSCRFL-5896).

## Supplementary Figure Legends

**Supplementary Figure 1.**
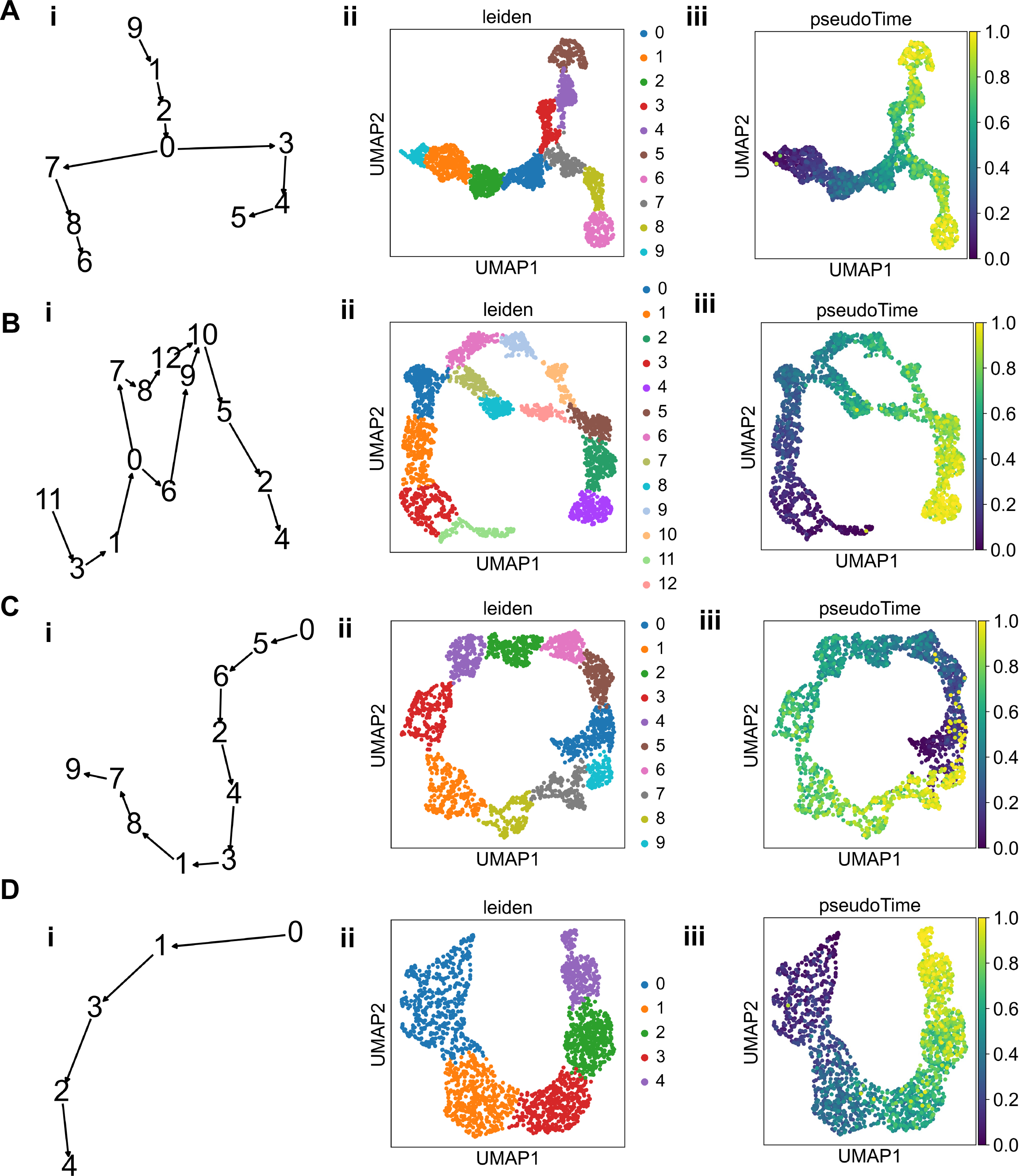
Manually curated input for BEELINE synthetic data that was used by OneSC for network inference benchmarking. (A) Dyn-BF data’s (i) cell state transition graph, (ii) cluster annotation overlayed on UMAP embeddings, (iii) inferred pseudotime overlayed on UMAP embeddings. (B) Dyn-BFC data’s (i) cell state transition graph, (ii) cluster annotation overlayed on UMAP embeddings, (iii) inferred pseudotime overlayed on UMAP embeddings. (C) Dyn-CY data’s (i) cell state transition graph that was broken up to form a linear structure, (ii) cluster annotation overlayed on UMAP embeddings, (iii) inferred pseudotime overlayed on UMAP embeddings. (D) Dyn-LI data’s (i) cell state transition graph, (ii) cluster annotation overlayed on UMAP embeddings, (iii) inferred pseudotime overlayed on UMAP embeddings.

**Supplementary Figure 2.**
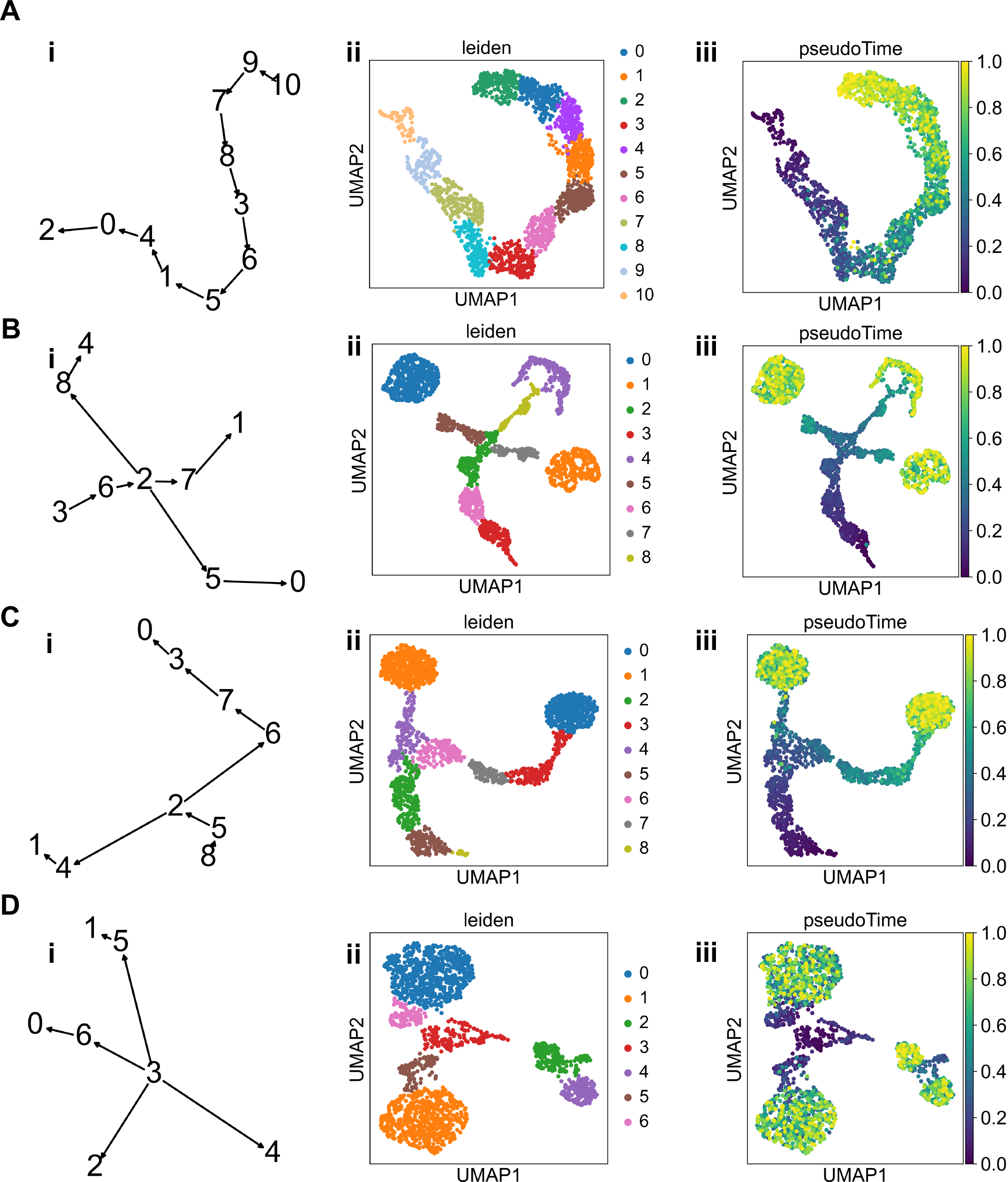
Manually curated input for BEELINE synthetic data that was used by OneSC for network inference benchmarking. (A) Dyn-LL data’s (i) cell state transition graph, (ii) cluster annotation overlayed on UMAP embeddings, (iii) inferred pseudotime overlayed on UMAP embeddings. (B) Dyn-TF data’s (i) cell state transition graph, (ii) cluster annotation overlayed on UMAP embeddings, (iii) inferred pseudotime overlayed on UMAP embeddings. (C) GSD data’s (i) cell state transition graph, (ii) cluster annotation overlayed on UMAP embeddings, (iii) inferred pseudotime overlayed on UMAP embeddings. (D) HSC data’s (i) cell state transition graph, (ii) cluster annotation overlayed on UMAP embeddings, (iii) inferred pseudotime overlayed on UMAP embeddings.

**Supplementary Figure 3.**
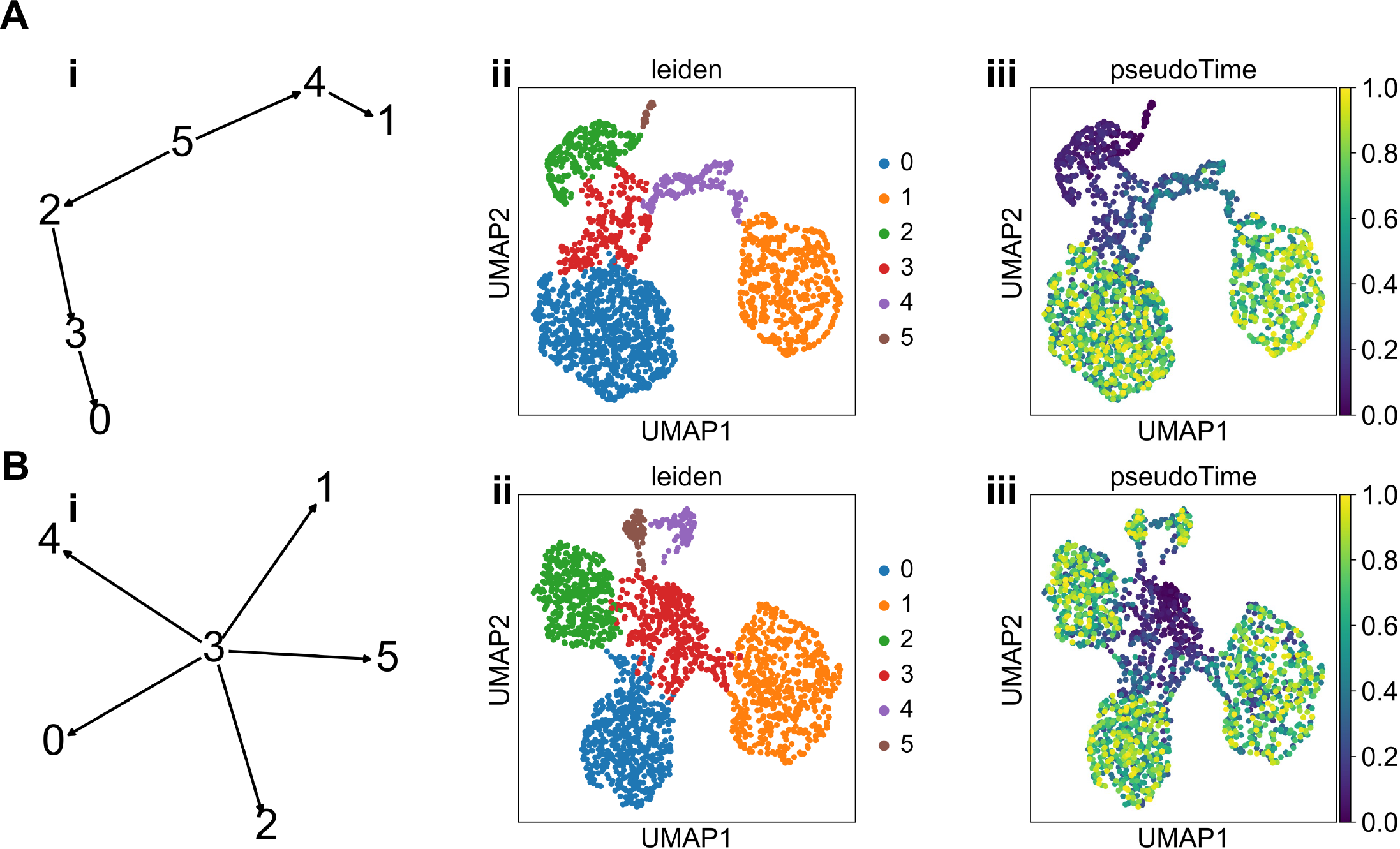
Manually curated input for BEELINE synthetic data that was used by OneSC for network inference benchmarking. (A) mCAD data’s (i) cell state transition graph, (ii) cluster annotation overlayed on UMAP embeddings, (iii) inferred pseudotime overlayed on UMAP embeddings. (B) VSC data’s (i) cell state transition graph, (ii) cluster annotation overlayed on UMAP embeddings, (iii) inferred pseudotime overlayed on UMAP embeddings.

**Supplementary Figure 4.**
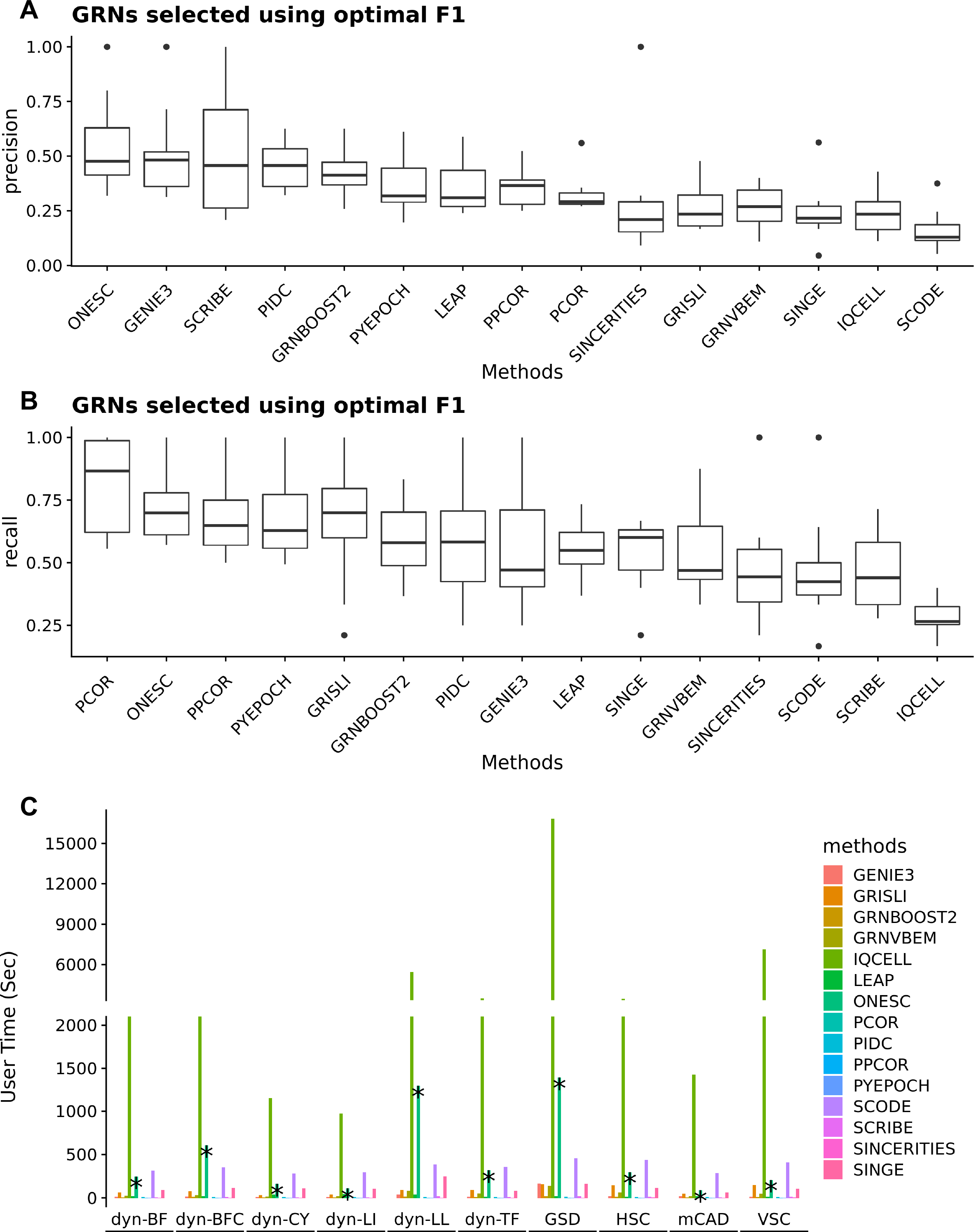
Benchmarking OneSC gene regulatory network inference with other methods using BEELINE platform. (A) Precision scores of F1 maximized networks from gene regulatory network inference methods tested on 10 BEELINE synthetic datasets (dyn-LI, dyn- LL, dyn-BF, dyn-BFC, dyn-TF, dyn-CY, mCAD, VSC, HSC, GSD). (B) Recall scores of F1 maximized networks from gene regulatory network inference methods tested on 10 BEELINE synthetic datasets. (C) Runtime of gene regulatory network inference methods tested on 10 BEELINE synthetic datasets. * indicates the runtime for OneSC.

**Supplementary Figure 5.**
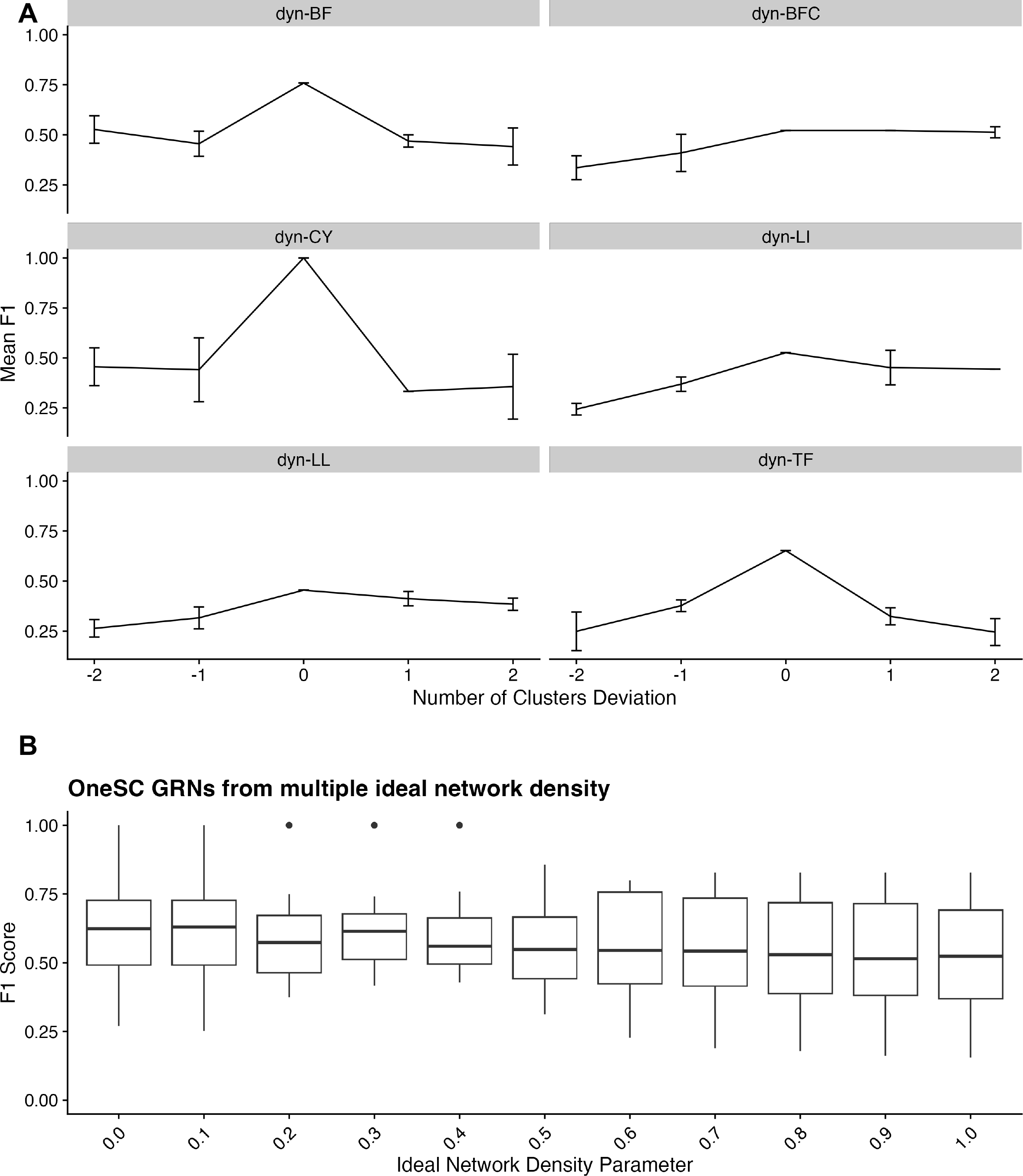
Parameter sensitivity testing of OneSC. (A) Line plots showing the change in mean and standard deviation of F1 scores when there is a deviation (-2, -1, +1, +2) in the cluster annotations through artificially splitting or merging clusters that were used to previously benchmark OneSC. There are 5 repeats per cluster deviation. (B) Box plot measuring the F1 score of OneSC network inferred using different ideal network densities.

**Supplementary Figure 6.**
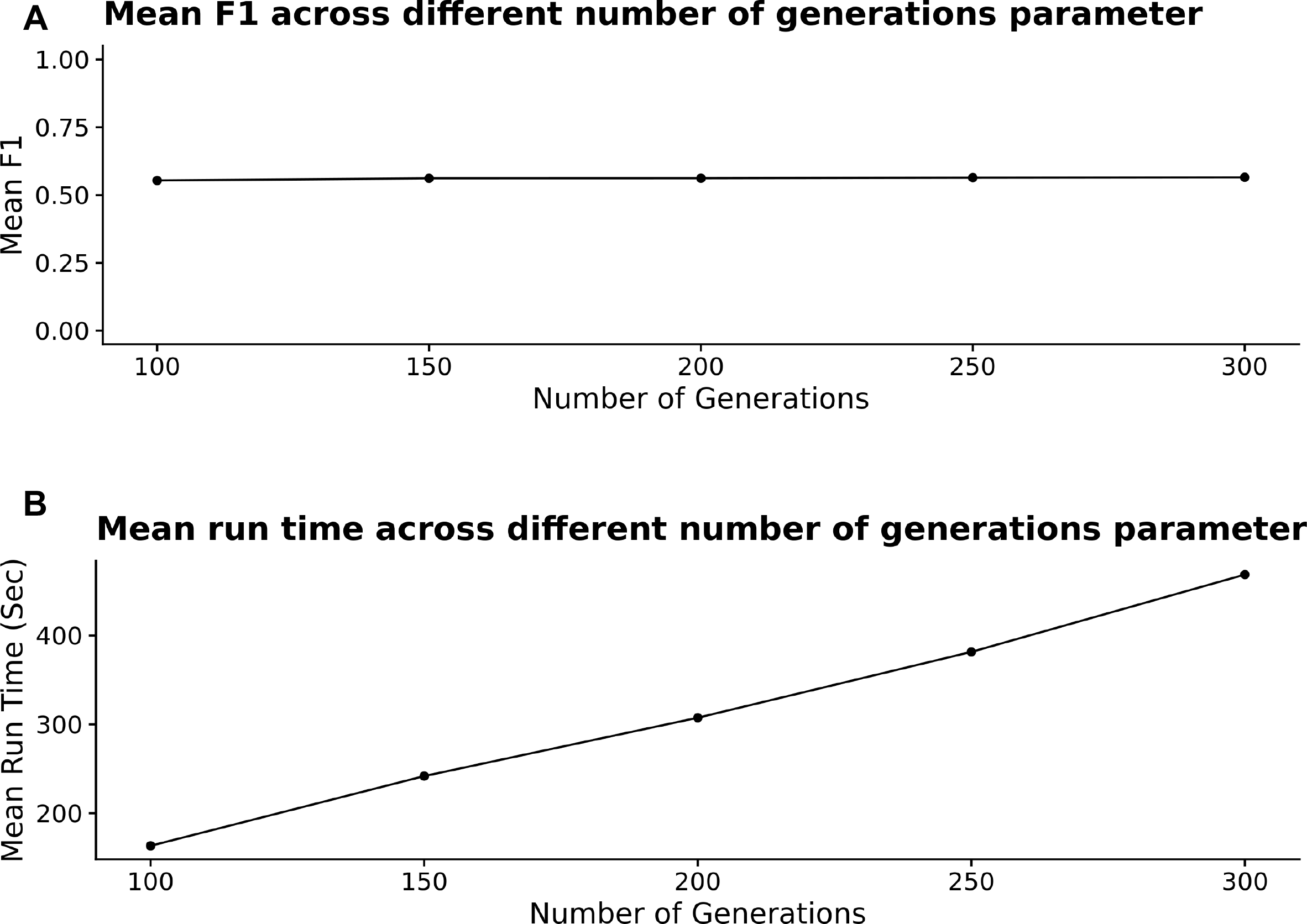
Testing the performance of OneSC using different number of generations for genetic algorithm. (A) Line plot showing the average F1 scores tested on 10 BEELINE synthetic datasets (dyn-LI, dyn-LL, dyn-BF, dyn-BFC, dyn-TF, dyn-CY, mCAD, VSC, HSC, GSD) using different parameters of genetic algorithm generations. (B) Line plot showing the average run-time tested on 10 BEELINE synthetic datasets using different parameters of genetic algorithm generations.

**Supplementary Figure 7.**
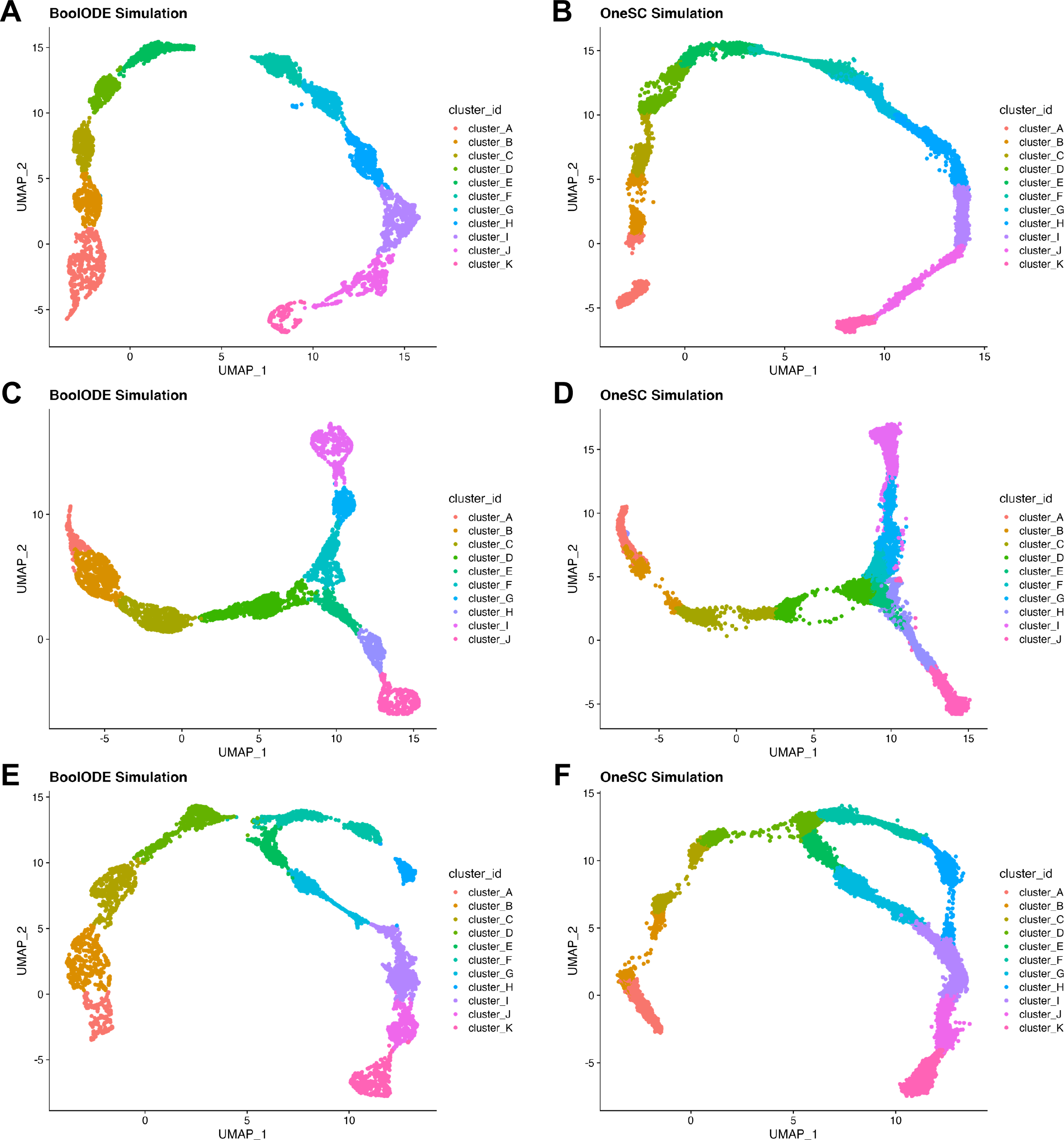
Comparisons of synthetic single-cell data simulated from OneSC and BoolODE. The cluster assignments for BoolODE synthetic data were done using standard Scanpy clustering. The cluster assignment for each cell in the OneSC synthetic data were labelled via finding the cell cluster with the highest Pearson correlation in expression profiles. UMAPs showing synthetic data generated from dyn-LI gold standard network by (A) BoolODE and (B) OneSC. UMAPs showing synthetic data generated from dyn-BF gold standard network by (C) BoolODE and (D) OneSC. UMAPs showing synthetic data generated from dyn-BFC gold standard network by (E) BoolODE and (F) OneSC.

**Supplementary Figure 8.**
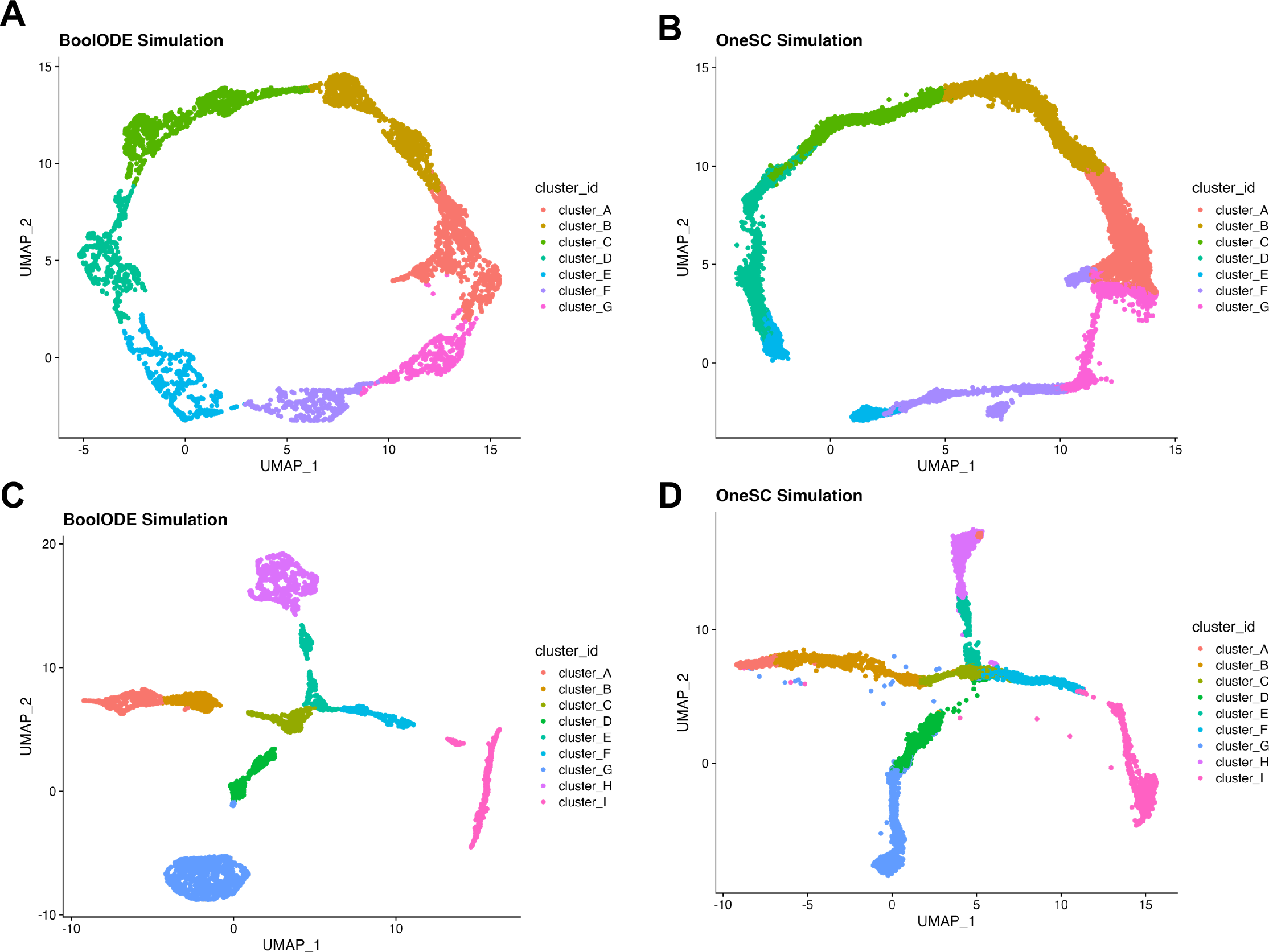
Comparisons of synthetic single-cell data simulated from OneSC and BoolODE. The cluster assignments for BoolODE synthetic data were done using standard Scanpy clustering. The cluster assignment for each cell in the OneSC synthetic data were labelled via finding the cell cluster with the highest Pearson correlation in expression profiles. UMAPs showing synthetic data generated from dyn-CY gold standard network by (A) BoolODE and (B) OneSC. UMAPs showing synthetic data generated from dyn-TF gold standard network by (C) BoolODE and (D) OneSC.

**Supplementary Figure 9.**
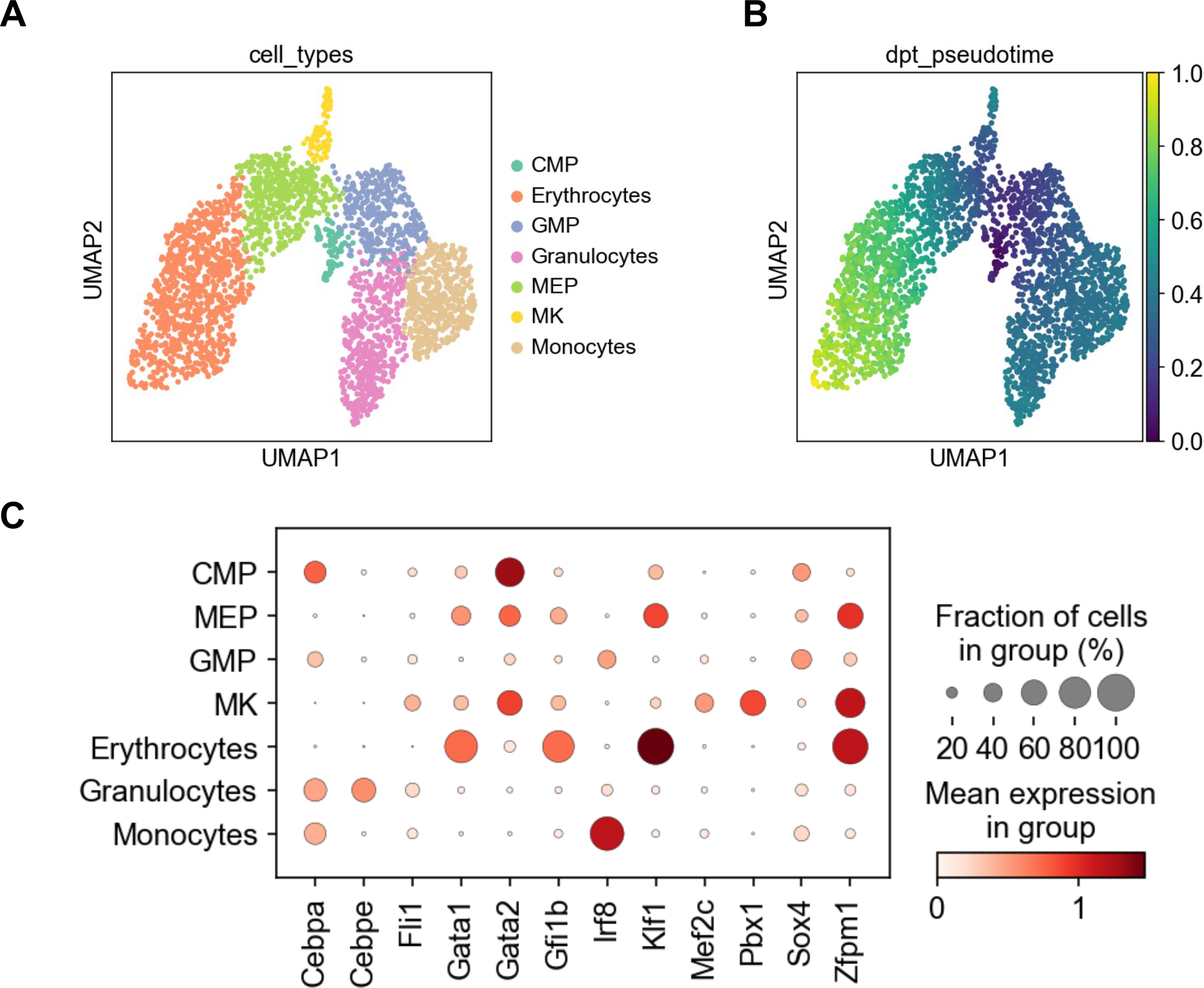
Preprocessing of Paul et al myeloid progenitor dataset. (A) UMAP showing the cell type cluster annotations. (B) UMAP showing the diffusion pseudotime assignments. (B) Dot plot showing the 12 dynamically expressed transcription factors across different cell types.

**Supplementary Figure 10.**
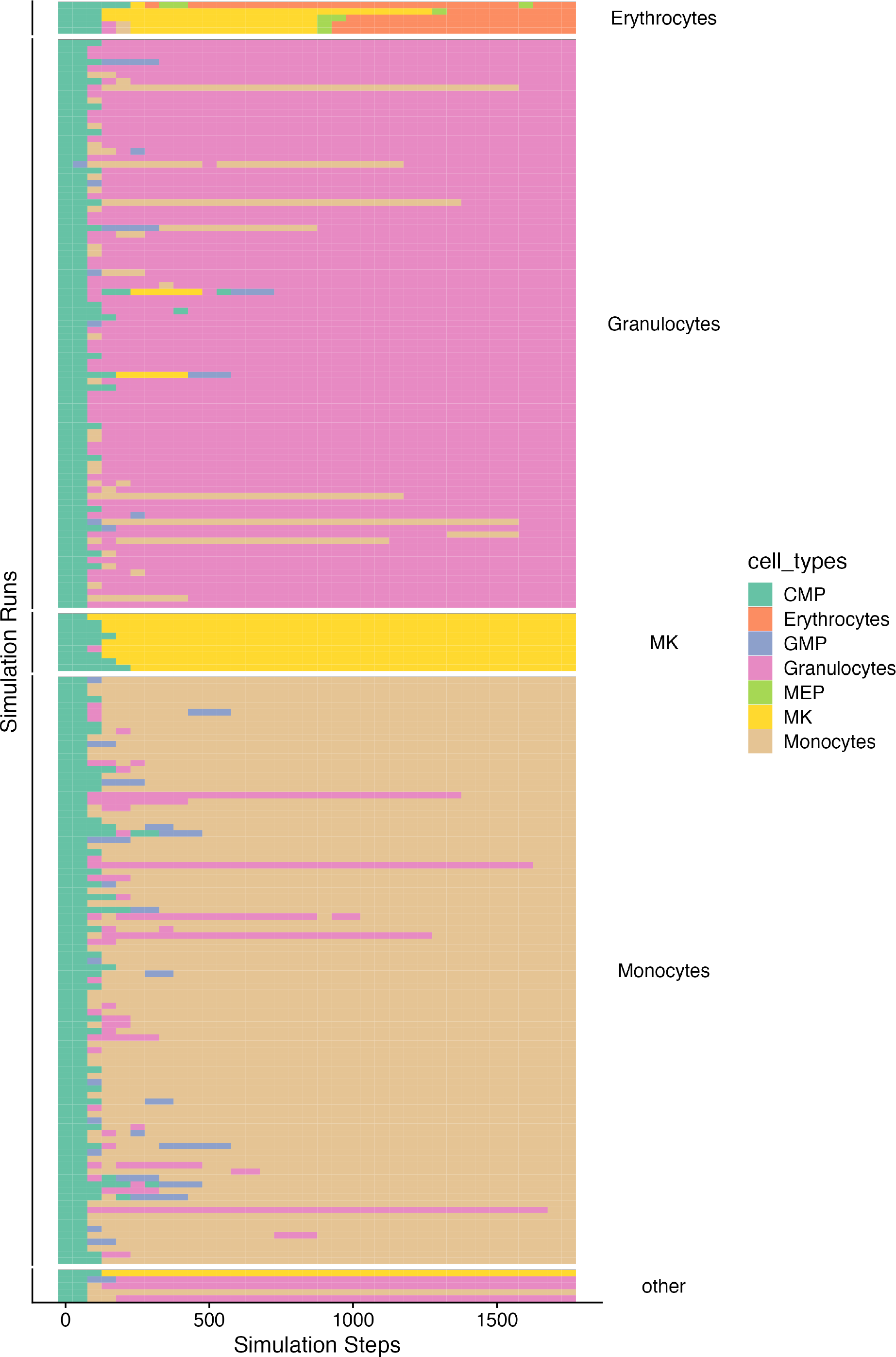
Heatmap showing the cell type assignment of 200 OneSC simulation runs from inferred network to capture myeloid differentiation trajectories. Each time step (a synthetic cell) in the simulation run is assigned to a cell type with the lowest distance in Boolean activity profiles.

**Supplementary Figure 11.**
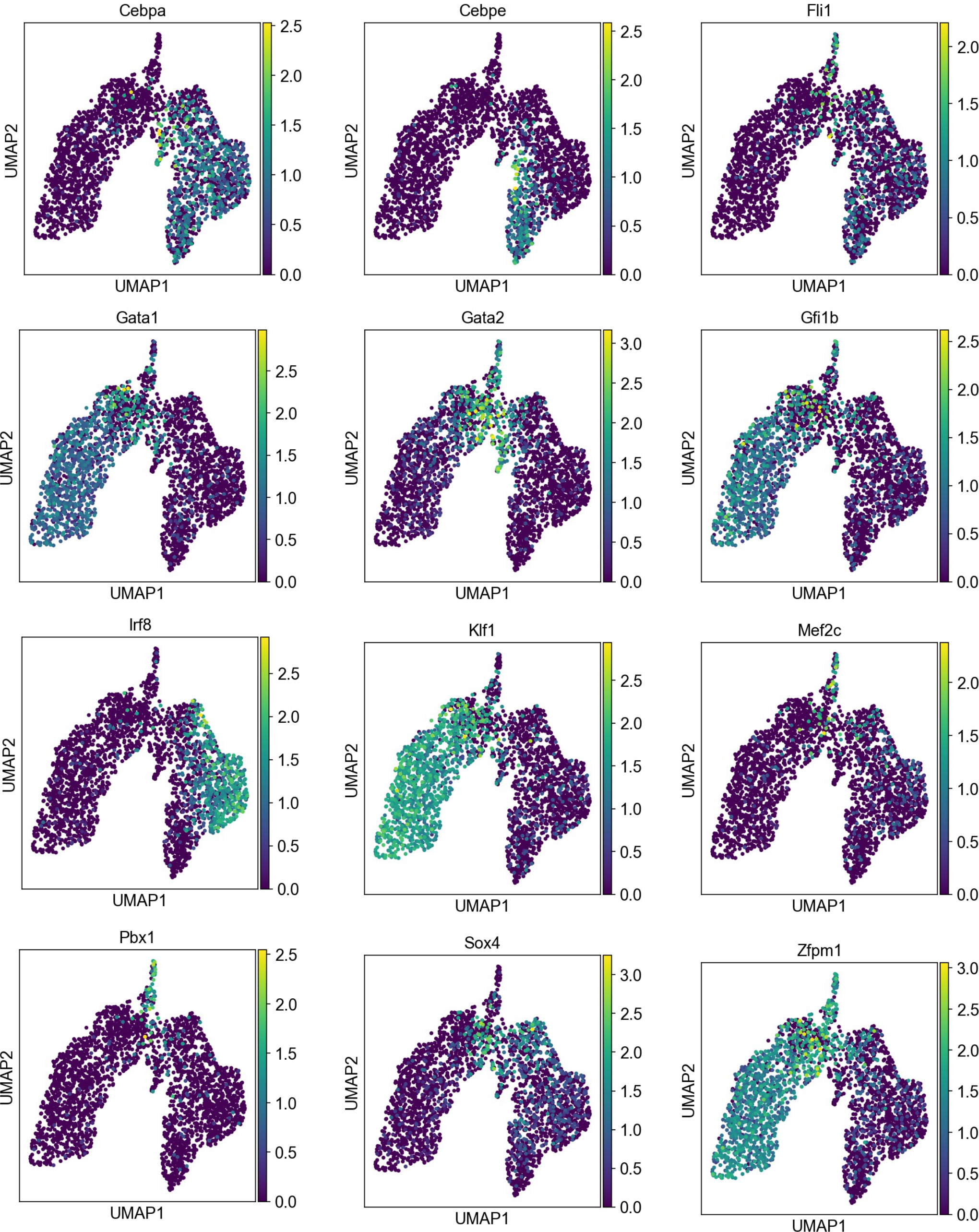
UMAPs showing the normalized expression values of the 12 dynamically expressed transcription factors in the Paul et al myeloid datasets.

**Supplementary Figure 12.**
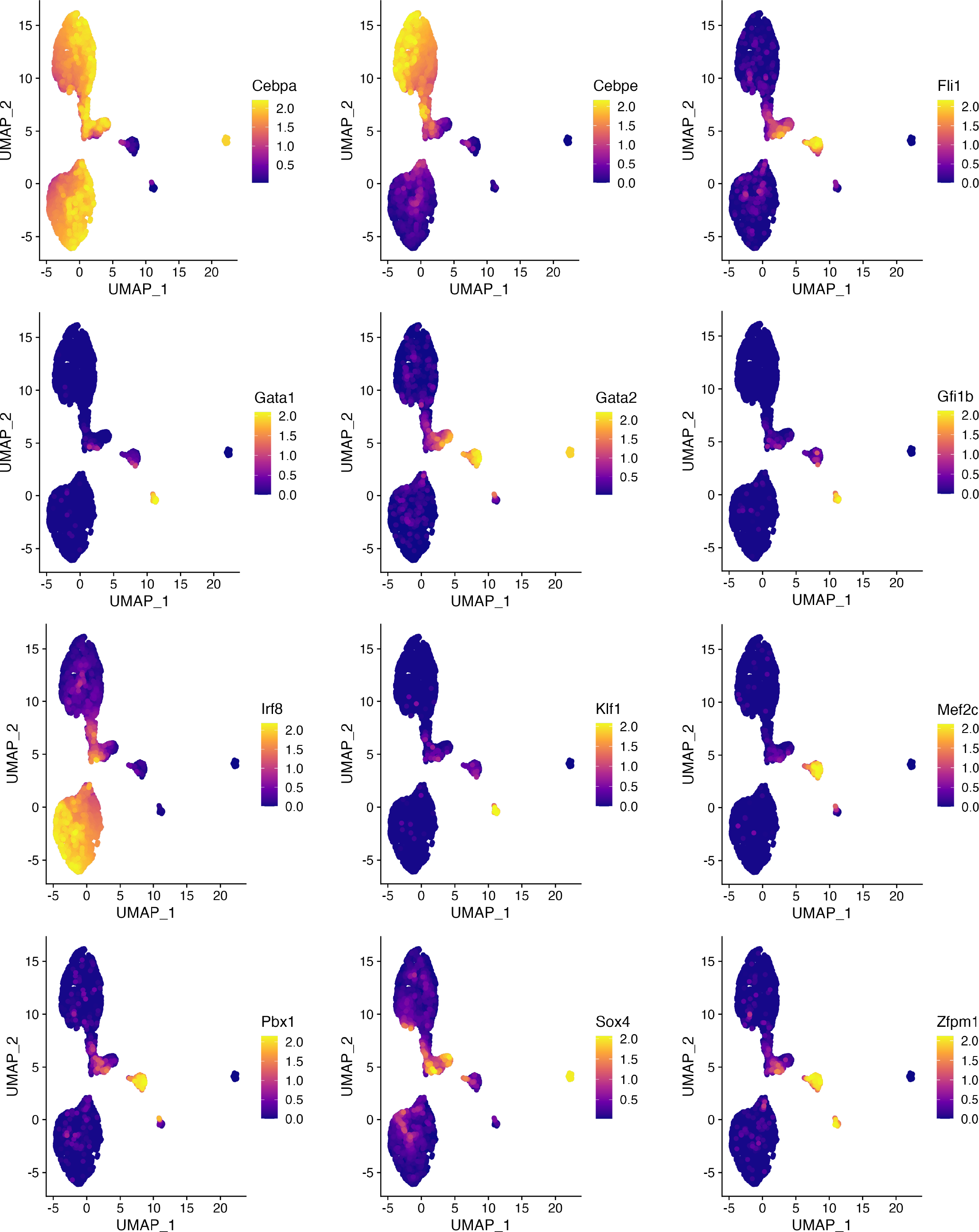
UMAPs showing the expression values of the 12 dynamically expressed transcription factors in the OneSC simulated cells.

**Supplementary Figure 13.**
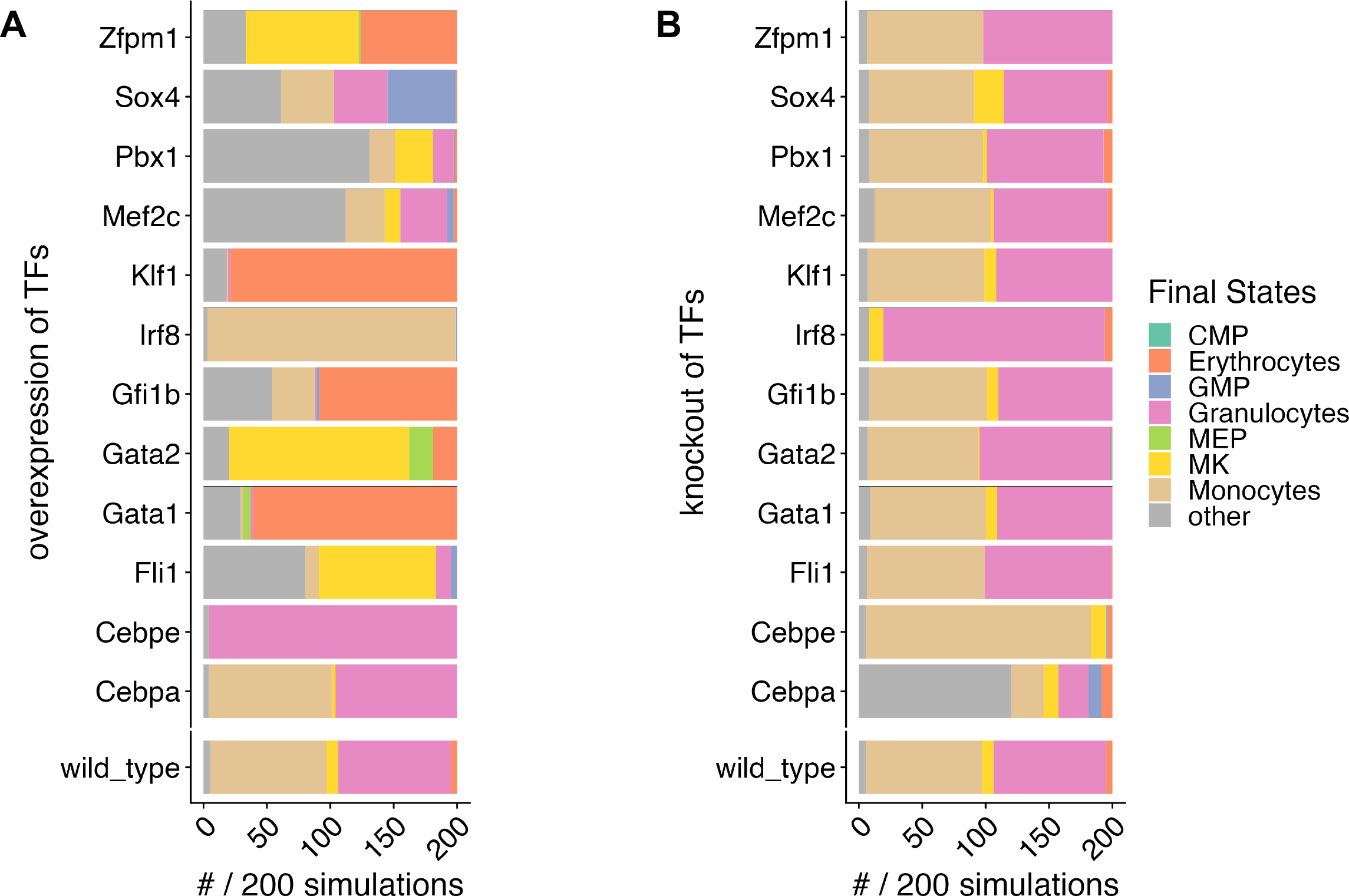
(A) Barplot showing the proportion of steady states from OneSC’s overexpression simulations of the 12 dynamically expressed TFs and no perturbation simulation (wild_type). (B) Barplot showing the proportion of steady states from OneSC’s knockout simulations of the 12 dynamically expressed TFs and no perturbation simulation (wild_type).

## Supplementary Table

**Supplementary Table 1.** Differentially expressed genes for cell type clusters in Paul et al dataset.

