## Supplementary Info for "OneSC: A computational platform for recapitulating cell state transitions"

**Supplementary Information**

1. **OneSC - Network inference method**
    A. **Construction of cluster transition graph**
    B. **Identification of dynamically expressed transcription factors:**
    C. **Curation of cluster/cell type level Boolean expression profiles**
    D. **Inference of executable GRNs using genetic algorithm**
2. **OneSC - Simulation method**
3. **Benchmarking GRN inference methods**
4. **Parameters stability test for OneSC**
5. **Comparisons of OneSC simulator and BoolODE**
6. **Processing of mouse myeloid progenitors single cell data**

### 1. OneSC – Network inference method:

The workflow of OneSC GRN inference is divided into four main sections A) construction of cluster transition graph, B) identification of the most dynamically expressed transcription factors, C) curation of cluster/cell type level Boolean expression profiles for regulatory network inference and D) network inference using genetic algorithm. Step 1 and 2 can be skipped if the transcription factors or cluster transition graph are defined independently.

#### A. Construction of cluster transition graph:

The initial step in the OneSC inference process involves creating a graph that illustrates the transitions between different cell type clusters along cell state transition trajectories. To do so, OneSC first calculates both the average expression profiles and the average pseudotime for each cell type cluster defined by the user. Then OneSC determines all feasible transitions from a "parent" cluster to a "child" cluster, with the condition that the average pseudotime of a parent must be less than that of its children. For each possible cell cluster transition pairs, a connectivity score is computed heuristically using the scaled Euclidean distance of the expression profiles and average pseudotime of parent cell cluster.

$$scaled\ dist_{AB} = \frac{dist_{AB} - dist_{min}}{dist_{max} - dis_{min}}$$

$$connectivity\ scores_{AB} = (scaled\ dist_{AB}) + (\frac{pseudotime_A}{2})$$

The rationale is that if cell type A is directly transitioning into cell type B, then the expression profiles should have the low distance between cell type A and cell type B and pseudotime of cell type A should be small.

For each cell type clusters that are not previously defined as initial or terminal (defined by user), OneSC identifies one incoming edge and one out going edge based on finding the lowest connectivity score among all the feasible transition pairs. For clusters that are defined as initial or terminal, OneSC will identify one incoming edge or one out going edge respectively based on finding the lowest connectivity score among all the possible transition pairs. Using this heuristic method, it is possible that two or more clusters can have incoming edges from the same parent cell cluster and consequently a parent cluster can also have multiple children clusters.

If the cell state transitions have been previously characterized, the users are free to manually curate the cell state transition graph using the networkx package.

#### B. Identification of dynamically expressed transcription factors:

Using user inputted cluster/cell type annotation and pseudotime ordering, OneSC identifies the most dynamically expressed transcription factors via modelling individual gene expressions across pseudotime using a generalized additive model (GAM) (Su *et al.*, 2022; Trapnell *et al.*, 2014). Users could define the adjusted p-value threshold (default 0.05), logfold change threshold (default 2) and minimal percent expression in cell type cluster (default 0.1). In the scenario of multiple trajectories, OneSC would run GAM modelling individually for each trajectory and gather all the TFs that are considered dynamically expressed in at least one trajectory.

If the core transcription factors that are previously characterized as important for differentiation in the biological system, the users can manually use those genes instead.

#### C. Curation of cluster/cell type level Boolean expression profiles:

Different from many scRNA-seq based GRN inference method, OneSC infers GRN based on the binarized cluster/cell type activity profiles rather than the expression profiles of individual cells. For each gene, OneSC curates the activity profiles (on/off) of itself as target gene labels and the activity profile for all the transcription factors in the network as the regulators profile across different cell states and different trajectories. The goal is to generate a subnetwork of regulators for each gene in the network such that the simulated Boolean output of the target gene based on regulators activity profiles is in concordance with target gene labels across different cell types. This section mainly focuses on the curation of target gene labels and regulators activity profiles of all the genes as input for genetic algorithm optimization. The details of the network inference and optimization process will be discussed in the next section.

OneSC first identifies the expression thresholds that binarize the gene expression profiles into 1 or 0 (on or off) for each gene. OneSC does so by averaging the genes expressions for each cell type cluster. Based on the highest averaged cluster expression ($clusterExp_{max}$) and the lowest average cluster expression ($clusterExp_{min}$), threshold for a gene is selected as

$$thresh = (clusterExp_{max} - cluster\mathrm{Exp}_{min})(percent_{cutoff}) + cluster\mathrm{Exp}_{min}$$

$percent_{cutoff}$ has a default value of 0.4 and can be adjusted by the users. A gene is considered turned on (activity status of 1) in a cluster if the gene expression is greater than the expression threshold and the cluster has at least 30 percent of the cells expressing the gene. Using the expression thresholds, OneSC generates the activity profiles of all genes across different cell states.

Next OneSC curates the activity status for each gene across all the cell states and their accompanying regulatory profiles. In cell states where the target gene does not change status from the previous cell state in the trajectory, the target gene status and regulators profile would just be the activity status of the genes at that cell state. If the target gene activity status does change at a cell state, then we have two scenarios at which the regulators profiles would have to be adjusted to reflect the subtle ordering at which different genes change during cell state transitions.

Using the inferred pseudotime, OneSC can determine the precise ordering at which genes are turned on or turned off to refine the inference of causal relationship between genes. Consider a situation where the activity status of a target gene changes at cell state N, and transcription factor A also changes activity status but after a regulatory window (default value set to 0.01 pseudotime unit prior to the target gene's activity shift), then activity status of transcription factor A at cell state N in the regulators activity profile for target gene would take the activity status of transcription factor A at the cell state N-1 because transcription factor A's activity status was still at cell state N-1 when target gene changed status. On the other hand, if target gene changes at cell state N but transcription factor A changes at an earlier point than target gene (before a regulatory window), then the activity status of transcription factor A at cell state N in the regulators activity profile for target gene would remain the status at cell state N.

To further use pseudotime ordering to finetune the causal relationship between genes, OneSC categorize transcription factors that are first turned on at a cell state later than target gene in a trajectory as ‘unlikely activator’. This is under the assumption that an activator should be turned on earlier than the target gene. During the network inference and optimization step, there will be a penalty term to discouraging transcription factor A from forming activating edge with target gene. However, it is possible to still infer transcription factor A as an activator if the activating edge increases the agreement between Boolean simulated status and observed target gene status. The specific details will be described in the next section.

OneSC repeats this data curation of target gene activity status and regulators activity profiles for all the genes in the network. If target gene is a transcription factor, it will also be a part of the regulators profile to potentially act as a self-activator. If the target gene is turned on at cell state N, then the activity status of target gene in the regulators profile for all the downstream cell states after cell state N are encoded as 1 (on). This is under the assumption that if the target gene can self-activate, once it is turned on it will be able to sustain itself.

To prioritize the reachability of steady states or final cell states in the development, the target gene status and regulators profile for terminal cell states are included twice in the training data. This way, the weight of reaching final cell states is increased during the network inference step. Lastly, cell states with conflicts of having same regulators profiles but different target gene status are removed as there are no possible network solutions.

#### D. Inference of executable GRNs using genetic algorithm:

With the target gene activity status labels and regulators profile of transcription factors for each gene, OneSC next optimizes the potential regulatory interactions (activation: 1, repression: -1, or no regulation: 0) between different regulators and target gene. OneSC uses a repressor centric scheme such that if there is at least one repressor active (activity status of 1), then the target gene activity is simulated to be inactive (activity status of 0). Thus, the Boolean rule of activators and repressors could be written as

$$(A_1 \vee A_2 \vee A_3 \vee \ldots) \wedge \neg R_1 \wedge \neg R_2 \wedge \neg R_3 \wedge \ldots$$

If at least one activator is active and no repressor is active then the target gene activity is active. If at least one activator is active and at least one repressor is active, then the target gene activity is inactive. If all activators are inactive then the target gene activity is inactive.

If there are no activators presented in regulatory network of target gene, then the Boolean rule is

$$\neg R_1 \wedge \neg R_2 \wedge \neg R_3 \wedge \ldots$$

If all repressors are inactive then the target gene activity is active. If at least one repressor is active then the target gene activity is inactive.

If there are no repressors presented in the regulatory network of target gene, then the Boolean rule is

$$A_1 \vee A_2 \vee A_3 \vee \ldots$$

If at least one activator is active then the target gene activity is active. If all activators are inactive then the target gene activity is inactive.

To perform optimization, OneSC first identifies the most optimal subnetworks for each gene and then combines all the subnetworks into a larger GRN. The target gene subnetwork is composed of regulators that have a direct edge towards the target gene. The subnetwork could be represented as a regulatory vector that represents the regulatory relationships of the regulators on the target gene with the length equal to the number of possible regulators.

$$\begin{bmatrix} v_1 \\ v_2 \\ v_3 \\ \ldots \\ v_{nreg} \end{bmatrix}$$

Each $v_i$ can be encoded in 3 ways $\{-1, 0, 1\}$ representing repression, no regulation and activation respectively for the target gene. Guided by the target gene activity status labels and regulator profiles across different cell states, OneSC uses genetic algorithm (Albadr *et al.*, 2020; Katoch *et al.*, 2021) to identify the regulatory relationship vector that maximizes the agreement between activity status of the target gene across different state outputted by the Boolean simulation (based on the activity status of assigned activators and repressors) and the activity status of target gene observed in the real data.

Genetic algorithm is a metaheuristic optimization process inspired by natural selection. In short, genetic algorithm first starts with an initial population of random subnetworks for a target gene. Each subnetwork is evaluated using a fitness function that favors agreement of the activity status of the target gene outputted by the Boolean logic and the activity status of target gene observed in the real data across different states. Then the set of subnetworks with high fitness are selected as parents and are subjected to crossover (random shuffling between subnetworks) and mutation (random changes to subnetworks) to generate next generation of subnetworks population. The process is repeated until it converges to an optimal fitness or reach the maximum generation limit. To overcome the stochastic nature of genetic algorithm and to have robust outputs, the process of subnetwork optimization is repeated multiple times with different initial populations and seeds for genetic algorithm. In the ensemble of inferred subnetworks, the majority subnetwork structure is voted to be optimal stable subnetwork. If there is a tie for the majority subnetwork, the subnetwork that has the number of edges closest to the ideal number of edges (set by the users) wins. OneSC uses PyGAD (Gad, 2021) (3.1.0) to perform genetic algorithm optimization.

The fitness function (higher values are better) for the genetic algorithm to maximize is made of several components that help achieve maximum target gene activity status agreement and also satisfy other secondary constraints. It is split into reward components (the higher the better) and penalty components (the lower the better).

$$
\begin{aligned}
fitness = {} & target\ gene\ correctness\ reward + additional\ edge\ reward \\
& + regulatory\ direction\ agreement\ reward + similar\ stage\ reward \\
& - unlikely\ regulator\ penalty - self\ inhibition\ penalty - no\ edge\ penalty \\
& - auto\ regulation\ penalty\ (can\ be\ turned\ off)
\end{aligned}
$$

$target\ gene\ correctness\ reward$ represents the reward given to the number of agreements in the activity status of target gene between Boolean simulation and activity status observed in training data across cell states. It is defined as

$$(\#\ of\ target\ gene\ activity\ status\ agreements)(10^4 \times n_{regulators})$$

The rationale for adding $n_{regulators}$ is so the reward scales with the number of genes in the network with the largest constant ($10^4$) such that achieving most amount of correct agreement for target gene activity status across cell states is the most important task and the center focus of the fitness function.

$additional\ edge\ reward$ represents the reward given to every new edge (activation or repression) added between regulators and target gene. The rationale is to promote adding new regulatory edges every new generation (if there are no other penalties). The reward is $10^2$ for each repressing or activating edge. However, if the number of regulator edges surpasses the ideal number of regulator edges (user defined parameter), then this term becomes $-10^2$ to promote the removal of unnecessary edges.

$regulatory\ direction\ agreement\ reward$ represents the reward given to every activating or repressing edge between regulators and target gene with the direction of regulation that match the direction of correlation between regulators and target gene. If the direction match, then the reward is $10 \times Pearson\ correlation$. Otherwise, the reward is $-10 \times Pearson\ correlation$. The rationale of this term is to promote regulatory interactions that match the sign of Pearson correlation.

$similar\ stage\ reward$ represents the reward given to regulatory edges in which regulators are first active within similar stage on the cell state transition hierarchy as the target gene. The assumption is that if a regulator is turned on around the same stage as the target gene, then it is more likely to be an activator or a mutual inhibitor (during cell fate decisions). All the regulators are assigned a weight that is inversely related to the cell state at which it is first active (i.e terminal states in the longest trajectory would get a weight of 1 and initial state would get a weight equivalent to the maximum number cell states along the longest trajectory). If regulator's weight is less or equal to target gene's weight + 1, then the reward would be $+\ regulator\ weight$ if there is an activating or repressing edge. This would allow the algorithm to prioritize regulators that are first active ideally one cell state earlier than the target gene. Otherwise, this term would be $-1$ to penalize scenarios where the regulator is first turned on much earlier than target gene.

$self\ inhibition\ penalty$ represents the penalty to prevent the formation of self-inhibition edge. The penalty is $3 \times 10^4 \times n_{regulators}$ which is 3 times the reward for matching the activity status of the target gene for a cell state. The rationale is to inhibit all potential auto-inhibitory edges. There are instances of self-repressing in transcription factors such as Hes familyn(Chen *et al.*, 2005; Bessho, 2003) resulting in oscillation in the expression dynamics, but there are insufficient amount of information from the mRNA expression profiles to properly infer this relationship as it typically requires additional sets of differential equations for protein dynamics to properly model the oscillation (Momiji and Monk, 2008).

$no\ edge\ penalty$ represents the penalty to prevent the situation of no regulatory edges for a target gene. The penalty is $3 \times 10^4 \times n_{regulators}$ which is 3 times the reward for matching the activity status of the target gene for a cell state. The rationale is to remove the possibility of having genes in the network that do not interact with any other genes or itself.

$unlikely\ regulator\ penalty$ represents the penalty for adding regulatory edges with regulators that were previously defined as unlikely to activate the target gene during the training data curation step. The penalty is $10^2$ per regulatory edge with unlikely regulator. By default, the unlikely activators are genes that are first active at a later cell state than the target gene. This penalty work in conjunction with $similar\ stage\ reward$ could help pinpoint the potential activators that are first active within a proximal window right before the target gene activation. OneSC also allows user to freely add unlikely activators or unlikely repressors for each target gene manually based on prior information.

$auto\ regulation\ penalty$ represents the penalty with every self-activating edge (could be turned off by the user). The rationale is to remove unnecessary self-activating edges. The penalty is $2 \times 10^2$.

**2. OneSC – Simulation method:**

OneSC, like BoolODE (Pratapa *et al.*, 2020), has a stochastic differential equation for each gene to model the dynamics of that gene's expression.

$$\frac{d[x_i]}{dt} = mf(Ri) - l_x[x_i] + s\sqrt{[x_i]}\,\Delta W_t$$

$$\Delta W_t = \mathcal{N}(0, h)$$

$[x_i]$ represents the mRNA expression level of target gene, $f(Ri)$ represents the regulatory function capturing all the regulators and their interactions with target gene, $m$ is the transcription rate (default value is 0.2), $l_x$ is the degradation rate (default value 0.1), $s$ is the strength of the noise (default value is 2), $\Delta W_t$ is the change in Wiener process increment sampled from a normal random distribution with mean zero and variance equivalent of simulation time step $h$ (default value is 0.01). OneSC use the Euler-Maruyama method (Kloeden and Platen, 1992) to numerically integrate the stochastic differential equations.

$$[x_i] = x_{i-1} + [mf(R_{i-1}) - l_x[x_{i-1}]](h) + s\sqrt{[x_{i-1}]}\,\Delta W_t$$

To perform in-silico perturbation, a perturbation term $D$ is added to the equation.

$$[x_i] = x_{i-1} + \left[(m + D)f(R_{i-1}) - l_x[x_{i-1}]\right](h) + s\sqrt{[x_{i-1}]}\,\Delta W_t$$

To simulate overexpression, assign $D$ to be a positive number (1 by default). To simulate knockout, assign $D$ to be a negative number (-1 by default). The range of expression for target gene during simulation is constrained to be between 0.02 and 2 by default if the expression values go outside of the range without the noise term.

We made two strategic decisions when implementing OneSC to enhance its computational efficiency. First, OneSC does not have equations to model protein dynamics. Second, instead of modelling all $2^N$combinations of TF states, OneSC employs Boolean algebra (Shannon, 1940) to model the transcription regulation. The algebraic forms of various Boolean logic are listed in Table 1.

| Boolean Operators | Boolean Algebra |
|---|---|
| $P_1$ or $P_2$ | $1 – (1-P_1)(1-P_2)$ |
| $P_1$ and $P_2$ | $(P_1)(P_2)$ |
| not $P_1$ | $1 – P_1$ |

Table 1. Boolean Operators and their corresponding Boolean algebraic form.

It is worth noting that similar approach was also used in the soft-heaviside version of BoolODE (Jalihal, 2020). $P$ in Table 1 represents the activity probability of a transcription factor scaling from 0 to 1 correlated with the expression values of the transcription factor. The activity probability is transformed from the expression values via a logistic function.

$$P_{TF} = \frac{1}{(1 + e^{-a([TF]-b)})}$$

$a$ adjusts for the steepness of the logistic function (default value is 7). $[TF]$ represents the expression values of the transcription factor. $b$ represents the mid-point between max and min expression values. Consider the example shown in (Pratapa *et al.*, 2020), if a gene (denoted by $X$) has two activators ($P, Q$) and one inhibitor ($R$)

$$X = (P \vee Q) \wedge \neg(R)$$

Combing Boolean polynomials, the regulatory function would then be written as

$$f(Ri) = \left(1 - \left(1 - P_p\right)\left(1 - P_Q\right)\right)(1 - P_R)$$

#### 3. Benchmarking GRN inference methods:

Synthetic data generated from BoolODE (Pratapa *et al.*, 2020) (network structures include LI, CY, LL, BF, BFC, TF, mCAD, VSC, HSC, GSD) were downloaded from https://doi.org/10.5281/zenodo.3378975 (version 3). For each synthetic gold standard network, the dataset (sample id 1) with 2000 cells and no dropout was used to benchmark OneSC against other methods. Clustering of the synthetic data was performed with Scanpy (1.9.1) (Wolf *et al.*, 2018) prior to running OneSC pipeline. 10 neighbors (n_neighbors=10) and 9 PCs (n_pcs=9) were used to construct the neighborhood graph (sc.pp.neighbors). The Leiden clustering resolution for the synthetic data were LI: 0.25 CY: 0.45, LL: 0.75, BF: 0.55, BFC: 0.55, TF: 0.3, mCAD: 0.15, VSC: 0.25, HSC: 0.25, GSD: 0.75. In the dyn-TF synthetic dataset, two clusters with similar expression states were combined forming now labelled cluster 4 (**Supp Fig 2B**). In the HSC synthetic dataset, two extremely small clusters (with fewer than 20 cells) were removed. All benchmarks using BEELINE platform (Pratapa *et al.*, 2020) were performed on AWS c5.4xlarge instance type. IQCELL was executed using default parameters shown in its tutorial.

To convert the edge weighted network into concrete networks for fair comparisons, a thresholding scheme was used to generate concrete networks that maximizes the F1 score. The precision, recall and F1 scores were calculated for each edge weight threshold. The edge weight threshold that produced the highest F1 score was chosen and all the edges with edge weights that were larger or equal to the threshold were kept as part of the concrete network. Different from how BEELINE calculates precision or recall, in order for a regulatory edge to be considered true positive, the edge must have the correct regulator, target gene, direction of regulation and type (activating or repressing). For methods that do not generate the type of regulation (GENIE3, GRISLI, GRNBOOST2, LEAP, PIDC, SCRIBE, SINGE), the sign of partial Pearson correlation was used.

Asynchronous Boolean update was used to simulate the steady states of inferred concrete networks using BoolNet R package (2.1.9) (Müssel *et al.*, 2010).

**4. Parameters stability test for OneSC:**

To explore the impact of over- and under-clustering, we randomly selected a cluster to either split (for over-clustering) or merge (for under-clustering). In the case of testing under-clustering, the selected cluster was then merged with its parent as defined by the cluster transition graph. When testing under-clustering, the random cluster was split in two such that cells with pseudotime < cluster mean pseudotime forme one cluster and all other cells are portioned into the second cluster. We tested the deviation of F1 scores as a result of over-clustering (+1 and +2 new clusters from optimal clustering) and under-clustering (-1 and -2 clusters from optimal clustering) on synthetic datasets dyn-BF, dyn-BFC, dyn-CY, dyn-LI, dyn-LL and dyn-TF. For each experimental condition, we repeated the process of over or under clustering 5 times.

The ideal_edges parameter represents the ideal number of regulators each target gene can have. The maximum number of regulators for a target gene is the total number of TFs in the network (including self-activation for target genes that are also TFs). The ideal network density parameter (ideal_edges parameter) was screened across different network density (0 to 1 with 0.1 increment) to test the change in F1 scores. The ideal_edges parameter was defined as

$$total\ \#\ of\ TFs \times\ network\ density$$

The number of generations parameter (num_generations) was screened across different values (100 to 300 with 50 increment) to test the change in F1 scores and run time.

**5. Comparisons of OneSC simulator and BoolODE:**

10 random networks were generated for each unique combinations of network sizes (5, 10, 15, 20) and network density (0.2, 0.4, 0.6, 0.8, 1) using networkx (Hagberg *et al.*, 2008) (version 2.8.8). BoolODE was cloned from https://github.com/Murali-group/BoolODE (commit id: 852b9b5d7e4aa95f9b37a301b551cd09d98b664a). To test the single-core performance, BoolODE simulations were done with no post processing (do_post_processing: false), no parallel (do_parallel: false), 1 core (nClusters: 1), 5 cell simulations (num_cells: 5) and simulation time of 50 (simulation_time: 50). The same was applied for soft-heaviside version of BoolODE. The above parameters gave 5000 distinct simulation steps for each cell simulation. OneSC simulations were ran 5 times with 5000 simulation steps for fair comparisons and runtime was recorded. To benchmark the simulators using parallel, the parallel option (do_parallel: true) is turned on and the number of cores is set to 5 (nclusters: 5) for BoolODE and the soft-heaviside version of BoolODE. OneSC simulations were ran with multiprocessing package using 5 cores. The benchmark was done on AWS c5.4xlarge machine.

To elucidate the similarity of synthetic cells generated from OneSC and generated from BoolODE using the same gold standard networks, we compared the cross correlation of gene dynamics and cluster similarity between synthetic cells generated from the two simulators. The simulation timesteps were scaled such that they are from zero to one, and cross-correlation of each gene was calculated for each distinct trajectory between OneSC synthetic cells and BoolODE synthetic cells across different gold standard networks (BF, BFC, CY, LI-extended and TF). While BF, BFC, CY were downloaded directly from https://doi.org/10.5281/zenodo.3378975 (5000 cells, sample id 1), we had to re-simulate LI structure again with simulation time of 7 instead of 5 for the simulated cells to fully reach the steady state. To compare cluster similarity, standard single-cell clustering was done on the synthetic dataset from BoolODE using Scanpy package (Wolf *et al.*, 2018), and the average expression profiles were generated for each cluster. Pearson correlations between each synthetic cell expression from OneSC and average expression profiles from BoolODE were generated. The synthetic cell from OneSC were labelled by the BoolODE category with the highest Pearson correlation.

**6. Processing of mouse myeloid progenitors single cell data:**

The mouse myeloid progenitors single-cell expression data were from Paul et al (Paul *et al.*, 2015) and downloaded using Scanpy's data loading function (scanpy.datasets.paul15()). Standard processing such as log-normalization, identifying highly variable genes (min_mean = 0.0125, max_mean = 3, min_disp = 0.5), scale expression (max_value = 10), neighborhood graph construction (n_neighbors = 10, n_pcs = 10), clustering (leiden cluster with resolution 0.5), diffusion pseudotime and cell type identifications were done using Scanpy package (Wolf *et al.*, 2018). The clusters were annotated based on the cluster marker genes. The synthetic cells were assigned to a cell type in the single-cell data by finding the highest correlation between synthetic cell's Boolean activity profiles and those of the cell types in the single cell data.
